# Mapping the genetic and environmental aetiology of autistic traits in Sweden and the United Kingdom

**DOI:** 10.1101/2020.04.14.031518

**Authors:** Zoe E. Reed, Henrik Larsson, Claire M.A. Haworth, Dheeraj Rai, Sebastian Lundström, Angelica Ronald, Abraham Reichenberg, Paul Lichtenstein, Oliver S.P. Davis

## Abstract

**Background:** Autistic traits are influenced by both genetic and environmental factors, and are known to vary geographically in prevalence. But to what extent does their aetiology also vary from place to place?

**Methods:** We applied a novel spatial approach to data on autistic traits from two large twin studies, the Child and Adolescent Twin Study in Sweden (CATSS; N=16,677, including 8,307 twin pairs) and the Twins Early Development Study in the UK (TEDS; N=11,594, including 5,796 twin pairs), to explore how the influence of nature and nurture on autistic traits varies from place to place.

**Results:** We present maps of gene- and environment-by geography interactions in Sweden and the United Kingdom (UK), showing geographical variation in both genetic and environmental influences across the two countries. In Sweden genetic influences appear higher in the far south and in a band running across the centre of the country. Environmental influences appear greatest in the south and north, with reduced environmental influence across the central band. In the UK genetic influences appear greater in the south, particularly in more central southern areas and the southeast, the Midlands and the north of England. Environmental influences appear greatest in the south and east of the UK, with less influence in the north and the west.

**Conclusions:** We hope this systematic approach to identifying aetiological interactions will inspire research to examine a wider range of previously unknown environmental influences on the aetiology of autistic traits. By doing so, we will gain greater understanding of how these environments draw out or mask genetic predisposition and interact with other environmental influences in the development of autistic traits.

## Introduction

Autism spectrum disorder (ASD) is a neurodevelopmental condition, manifesting in childhood and characterised by persistent difficulties with social communication and repetitive behaviours. ASD has a significant impact on child development, often including language difficulties and other co-occurring conditions which may persist into adulthood (Seltzer, Shattuck, Abbeduto, & Greenberg, 2004).

The aetiology of ASD reflects both genetic and environmental influences. Twin and family studies suggest that genetic differences between people explain around 80% of the population variance for ASD (Bai et al., 2019; Sandin et al., 2017; Tick, Bolton, Happé, Rutter, & Rijsdijk, 2016). Most studies suggest that the remaining variance is explained by non-shared environmental influences — that is, environmental influences that do not contribute to greater similarity within families.

Reported prevalence of ASD in developed countries varies between 1% and 3%, with higher estimates in recent years (Delobel-Ayoub et al., 2020; Idring et al., 2015; Lundström, Reichenberg, Anckarsäter, Lichtenstein, & Gillberg, 2015; Lyall et al., 2017; Maenner et al., 2020), although this is known to vary across geographical regions (Bakian, Bilder, Coon, & McMahon, 2015; Campbell, Reynolds, Cunningham, Minnis, & Gillberg, 2011; Chen, Liu, Su, Huang, & Lin, 2008; Delobel-Ayoub et al., 2020; Hoffman et al., 2017; Mazumdar, Winter, Liu, & Bearman, 2013). Several studies suggest that living in an urban environment is associated with greater risk of ASD compared to rural environments (Chen et al., 2008; Lauritsen et al., 2014; Vassos, Agerbo, Mors, & Bøcker Pedersen, 2016; Wu & Jackson, 2017). Possible reasons given for this geographical variation in prevalence of ASD include regional diagnostic bias, differences in access to health services or diagnostic resources (Chen et al., 2008; Lauritsen et al., 2014; Vassos et al., 2016), different levels of parental awareness (Lauritsen et al., 2014; Vassos et al., 2016), air pollution exposure during pregnancy (Weisskopf, Kioumourtzoglou, & Roberts, 2015), green space availability (Wu & Jackson, 2017) and local trends in socioeconomic status (Bakian et al., 2015; Lyall et al., 2017).

If prevalence of autistic traits varies from place to place, is the same true of the aetiology? For example, does variation in the environment explain variation in autistic traits in some areas more than others? Or does the environment in some areas draw out genetic differences between children in their propensity for developing autistic traits? We previously developed a spatial approach to twin model-fitting called spACE to detect spatial variation in genetic and environmental influences within a country (Davis, Haworth, Lewis, & Plomin, 2012). This approach has the potential to highlight gene-environment and environment-environment interactions for outcomes such as autistic traits. These interactions represent variation in aetiological influences on a trait depending on environmental exposure. For example, genetic risk of a mental health disorder may be drawn out by a stressful environment or genetic risk of hay fever may only reveal itself in pollen-rich areas. The spACE approach allows us to investigate this, mapping geographical patterns of nature and nurture without requiring the measurement of specific genetic variants or specific environmental characteristics.

Here we apply the spACE approach to data on autistic traits in Sweden and the UK. Autistic traits and diagnostic categories of ASD show substantial aetiological overlap (Colvert et al., 2015; Robinson et al., 2016), with genetic correlations from bivariate twin models of 0.52-0.89. The heritability of autistic traits does not change as a function of severity (Lundström et al., 2012; Robinson et al., 2011; Ronald et al., 2006), and genetic links have been identified between extreme and sub-threshold variation in ASD (Robinson et al., 2011; Ronald et al., 2006), so to maximise power we have focussed on trait measures rather than diagnoses.

We hope that by systematically mapping geographical differences in aetiology we will facilitate identification of new environments and shed light on the mechanisms by which they act.

## Methods and materials

### The Swedish Twin Registry and CATSS

The Child and Adolescent Twin Study in Sweden (CATSS) (Anckarsäter et al., 2011), a sub-study of the Swedish twin registry (Magnusson et al., 2012), was launched in 2004 to investigate childhood-onset neurodevelopmental problems such as ADHD and ASD in childhood and adolescence, for all twins turning 9 or 12 years since 2004. Parents were asked to participate in a telephone interview to collect information on various health-related issues. By 2013, when data on autistic traits were obtained, 8,610 parents had responded to this request, accounting for 17,220 twins. In 87.5% of these interviews, the informant was the mother (Anckarsäter et al., 2011). The CATSS-9/12 study obtained ethical approval from the Karolinska Institute Ethical Review Board: Dnr 03-672 and 2010/507-31/1, CATSS-9 - clinical 2010/1099-31/3 CATSS-15 Dnr: 2009/1599-32/5, CATSS-15/DOGSS Dnr: 03-672 and 2010/1356/31/1, and CATSS-18 Dnr: 2010/1410/31/1.

For autistic traits, 16,677 participants had data available (including 8,307 complete pairs and 63 incomplete pairs of twins).

### CATSS measures of autistic traits

The Autism-Tics, ADHD and other Comorbidities (A-TAC) inventory, based on the Diagnostic and Statistical Manual of Mental Disorders (DSM)-IV criteria, was used in the telephone interview with parents to collect information on a range of neurodevelopmental problems. This inventory has previously been validated in both clinically diagnosed children and the general population (Anckarsäter et al., 2008; Hansson et al., 2005; Larson et al., 2010, 2013; Mårland et al., 2017) and reliability, as measured by Cronbach’s alpha, showed high internal consistency (Cronbach’s alpha = 0.86), with further details previously reported (Anckarsäter et al., 2011). The inventory includes 17 items that assess autistic traits, where respondents can answer ‘yes/1’, ‘yes, to some extent/0.5’, and ‘no/0’. Following the standard approach, we created a score for each individual by summing these item scores. In previous validation studies a low and high cut-off of 4.5 and 8.5 for ASD have been established for broad screening and for use as a clinical proxy, respectively. We standardised the phenotype data to mean 0 and SD 1 at the population level to simplify comparisons between the UK and Sweden and to make it easier to identify areas where the total variance is greater than or less than the population average.

### CATSS location data

To conduct the spACE analysis, we assigned a geographical location to each family. In CATSS we matched each twin pair to a Small Areas for Market Statistics (SAMS) location, for the most recent location data we had available up to 2009, using data from Statistics Sweden (http://www.scb.se/en/) and assigned coordinates based on the centroid of the SAMS location.

To provide context for the results for Sweden, it is useful to understand a little about its geography. **Figure 1** shows a map of Sweden and some general indicators of the country’s geography; the **supplementary materials** contain a detailed description.

**Figure 1.**
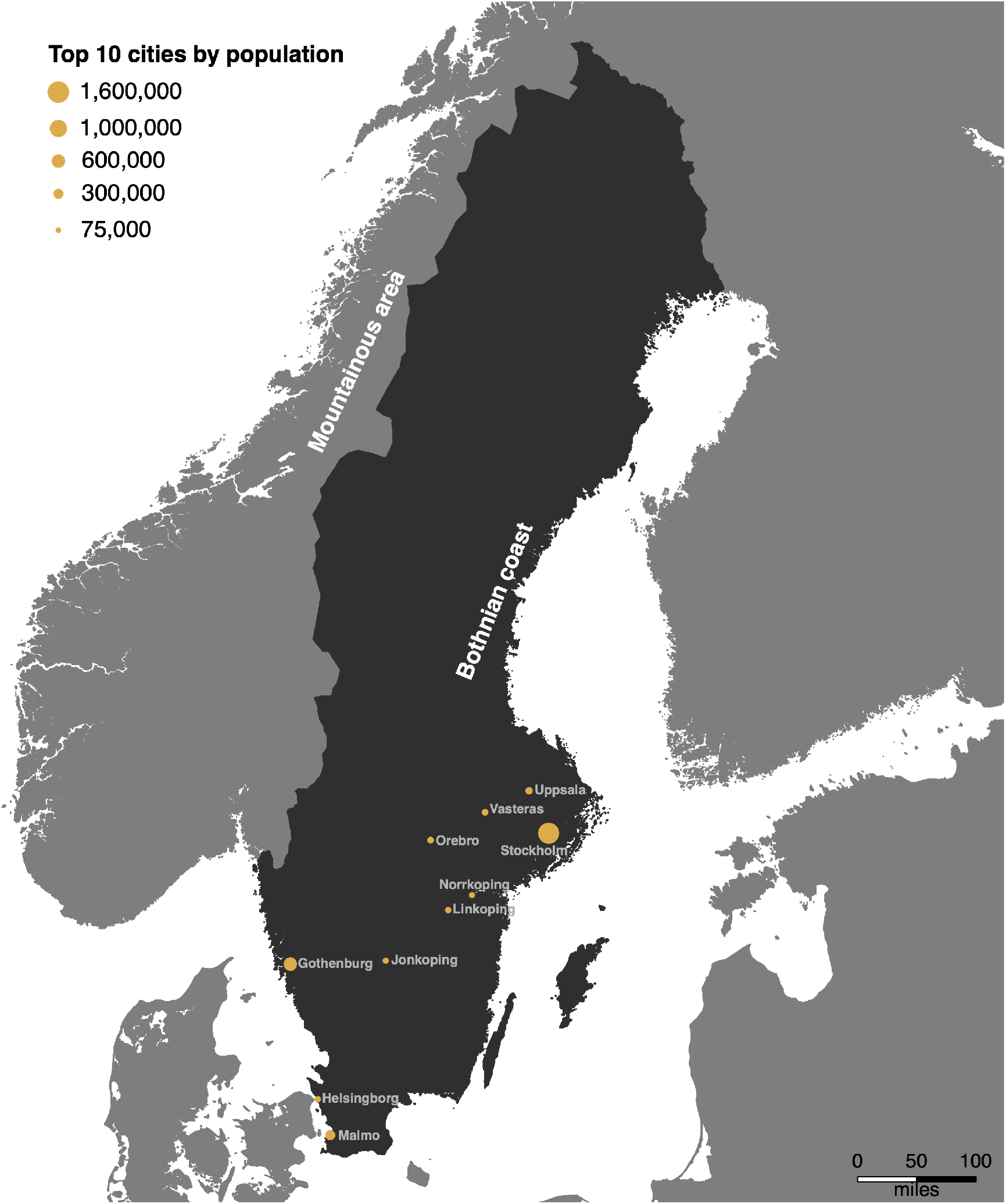
Map of Sweden with top 10 cities by population. Sweden is shown in dark grey with the surrounding countries in a lighter grey. The top 10 most populated cities are indicated by orange circles, where the area reflects the population.

### The Twins Early Development Study

The Twins Early Development Study (TEDS) contacted parents of twins born in England and Wales between January 1994 and December 1996 (Haworth, Davis, & Plomin, 2013). 16,810 pairs of twins were initially recruited, and currently there are over 10,000 twin pairs still enrolled in TEDS. The participants are demographically representative of the UK population of a similar age, with the majority identifying themselves as white British and with English as their first language. TEDS has collected wide-ranging data on cognitive and behavioural development, using approaches that include questionnaire booklets, telephone testing and web-based tests. The twins, their parents and teachers have all participated in data collection. However, there are no data available on which parent completed the questionnaire for this study. Ethical approval for TEDS research is provided by the Institute of Psychiatry, Psychology and Neuroscience Ethics Committee, King’s College London.

Full phenotypic data for autistic traits were available for 11,594 TEDS participants (including 5,796 complete pairs and 62 incomplete pairs of twins).

### TEDS measures of autistic traits

Parents in TEDS completed the Childhood Autism Spectrum Test (CAST) when the twins were age 12 years. Despite some attrition, the sample who provided CAST data at age 12 remain reasonably representative of the UK population when compared to demographic data from the UK’s Office for National Statistics (Robinson et al., 2012). The CAST consists of 30 items, scored 1 for yes or 0 for no (Scott, Baron-Cohen, Bolton, & Brayne, 2002). The CAST has been shown to have good internal consistency (Cronbach’s alpha = 0.71 to 0.81, depending on age and rater) (Holmboe et al., 2014), good test-retest reliability with a correlation of 0.83 between tests (J. Williams et al., 2006). The CAST score considered indicative of ASD is 15. Once again, we standardised the phenotype data to mean 0 and SD 1 at the population level to simplify comparisons between the UK and Sweden and to make it easier to identify areas where the total variance is greater than or less than the population average.

### TEDS location data

We assigned each twin pair geographical coordinates based on the centroids of their postcodes at age 12. To provide context for the results for the UK, **Figure 2** displays a map and some general indicators of the country’s geography; the **supplementary materials** include a detailed description.

**Figure 2.**
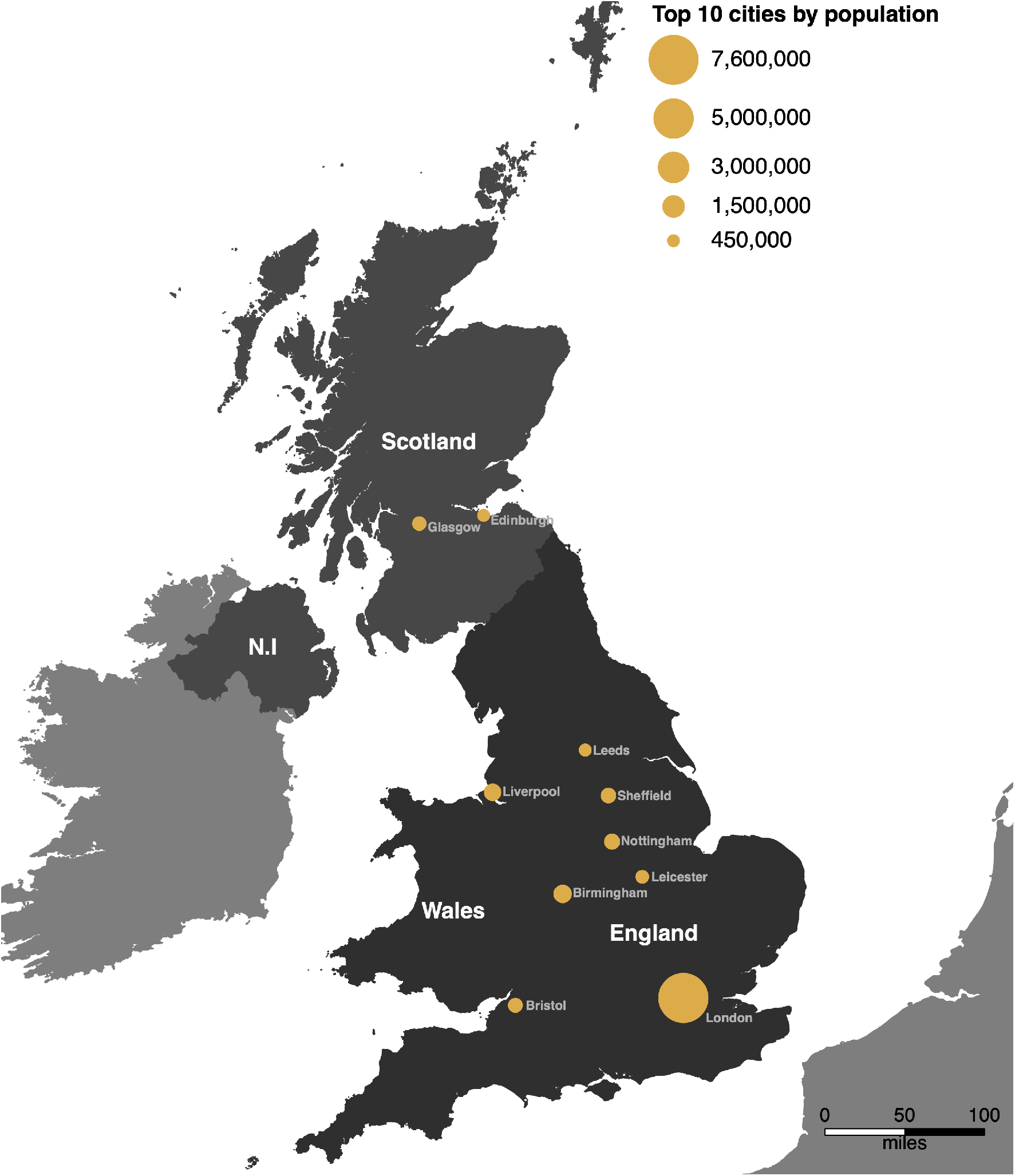
Map of the UK with top 10 cities by population. The recruitment area (England and Wales) for the Twins Early Development Study (TEDS) is shown in dark grey, with the rest of the UK (Scotland and Northern Ireland [N.I]) in a lighter grey. Other countries are shown in the lightest shade of grey. The top 10 most populated cities are indicated by orange circles, where the area reflects the population.

### Statistical analyses

#### ACE models and maps in CATSS and TEDS

In twin analysis, within-pair similarity of monozygotic (MZ) and dizygotic (DZ) twins is compared to estimate parameters for additive genetic (A), shared environmental (C) and non-shared environmental (E) influences on a trait. In this context, the shared environment refers to influences other than DNA similarity that make children growing up in the same family more similar to each other, whilst the non-shared environment refers to influences that do not contribute to similarity within families and also includes error in measuring the phenotype. It is not possible to assign specific environments to one or the other environmental component because most environments themselves show both shared and non-shared (and often genetic) influences. Although it is tempting to assign specific events or circumstances such as parental divorce or sports injury to one or the other, in most cases the effects of specific events or circumstances is complex and difficult to assign. For example, divorce is often experienced differently by children in the same family, and the propensity to receive sports injuries is partly genetically influenced. We can estimate the contribution of genetic and environmental influences because of the different ways these influences are shared in MZ and DZ twin pairs. For MZ twins, who share 100% of their segregating alleles, A influences correlate 1, whereas for DZ twins they correlate 0.5 because DZ twins share, on average, 50% of their segregating alleles. For both MZ and DZ twins growing up in the same family the shared environmental correlation is 1. In contrast, the non-shared environment is uncorrelated and contributes to differences between twins (Rijsdijk & Sham, 2002).

In this study, we applied a version of the spACE analysis method (described in detail and validated in (Davis et al., 2012)) to explore how A, C and E for autistic traits vary geographically. We fit full information maximum likelihood structural equation models to twin data in R (version 3.3.1) using the OpenMx package (version 2.9.4), calculating A, C and E at many different target locations across an area. In this study we built on our previous work by applying the weights within the structural equation modelling framework, rather than by calculating weighted correlation matrices and using those as input. In twin analysis it is possible to model non-additive genetic effects (D) instead of shared environmental effects (C). D influences are sometimes found with ASD (Holmboe et al., 2014), although often the D component can be dropped in favour of an AE model (Taylor, Charman, & Ronald, 2015; Taylor, Gillberg, Lichtenstein, & Lundström, 2017).

However, the D component is highly correlated with the A component, which means confidence intervals are wide and the tendency of variance to swap between these two components makes it difficult to compare results across locations. In other words, if we were to include a D component this is likely to introduce noise into our results due to the high correlation between A and D. Because of this, we have not modelled D here in our main analyses but have included ADE maps in the supplementary materials. This means that the A component, which usually represents additive genetic influences, will also absorb any non-additive genetic influences.

For target locations in Sweden we used the centroid of each unique SAMS that included at least one twin pair. Because UK postcodes give more precise locations than Swedish SAMS, we instead selected UK target locations representative of local population density to preserve participant anonymity. We selected target locations in the UK and Sweden to reflect population density. However, the choice of target locations for plotting does not affect the results at all and any other collection of locations could equally be used. All twin pairs contributed to the results at each location, but contributions were weighted according to the distance of each twin pair from the target location:

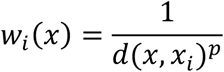

where *x* represents the target location, *x_i_* represents the location of a twin pair, *d* is the Euclidean distance between *x* and *x_i_*, and *p* is the power parameter that controls the rate of drop-off of a twin pair’s influence over distance (0.5 for these analyses). This tuning parameter was selected as a trade-off between accurately estimating A, C and E components and accurately localising them. The value we selected results in some smoothing of estimates towards population means, but still allows patterns of variation to be seen. We included sex as a covariate in all the models (accounting for 2.59% of the variance), and age in the TEDS data (accounting for 0.34% of the variance). Further detail on the spACE approach can be found in (Davis et al., 2012). We plotted maps to visualise results (**figures 3 and 4**), where each target location is coloured according to the value of the estimate at that location compared to the mean of values across the map. Low values appear blue and high values appear red, with increasing salience of the colour representing increasing distance from the mean. To avoid outliers having a large effect on the distribution of colours in the maps, we assigned the highest 4% of values to the brightest red and the lowest 4% of values to the brightest blue before assigning colour values to equal ranges between the two. The histograms show the distribution of results and the corresponding colours. Higher resolution and interactive versions of the maps can be found at https://dynamicgenetics.github.io/spACEjs/.

**Figure 3.**
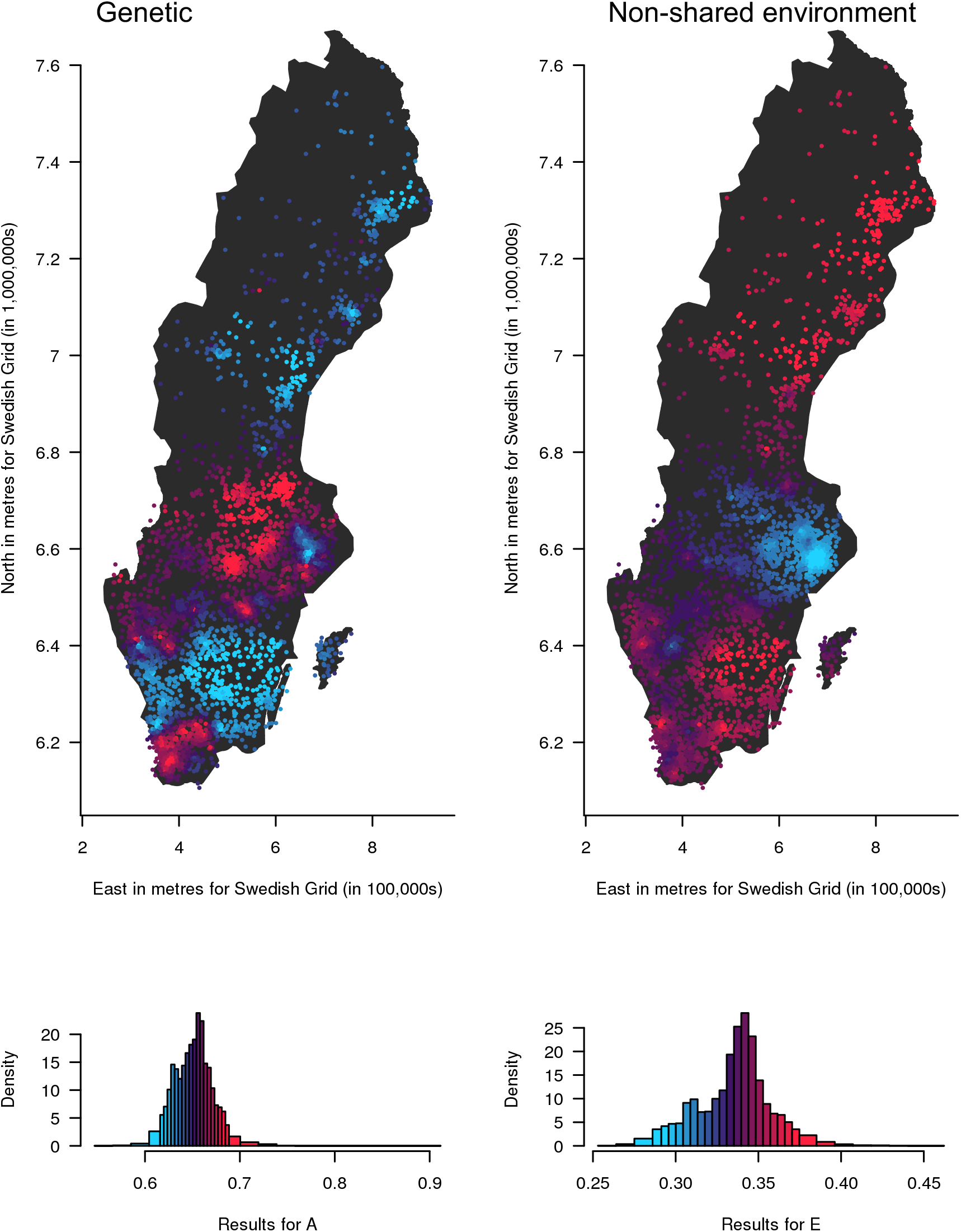
Mapping genetic (A) and non-shared environmental (E) influences on autistic traits in Sweden shows that both vary geographically across the country. Geographical variation in genetic (A) and non-shared (E) influences on childhood autistic traits in Sweden (results overlaid on an outline of the SAMS areas). The contributions of A and E range from low (blue) to high (red). The histograms show the distribution of the estimates, coloured in the same way as the points on the map. The estimates are not standardised and are therefore not constrained to add up to one. A higher-resolution interactive version of this map is available at https://dynamicaenetics.aithub.io/spACEjs/.

**Figure 4.**
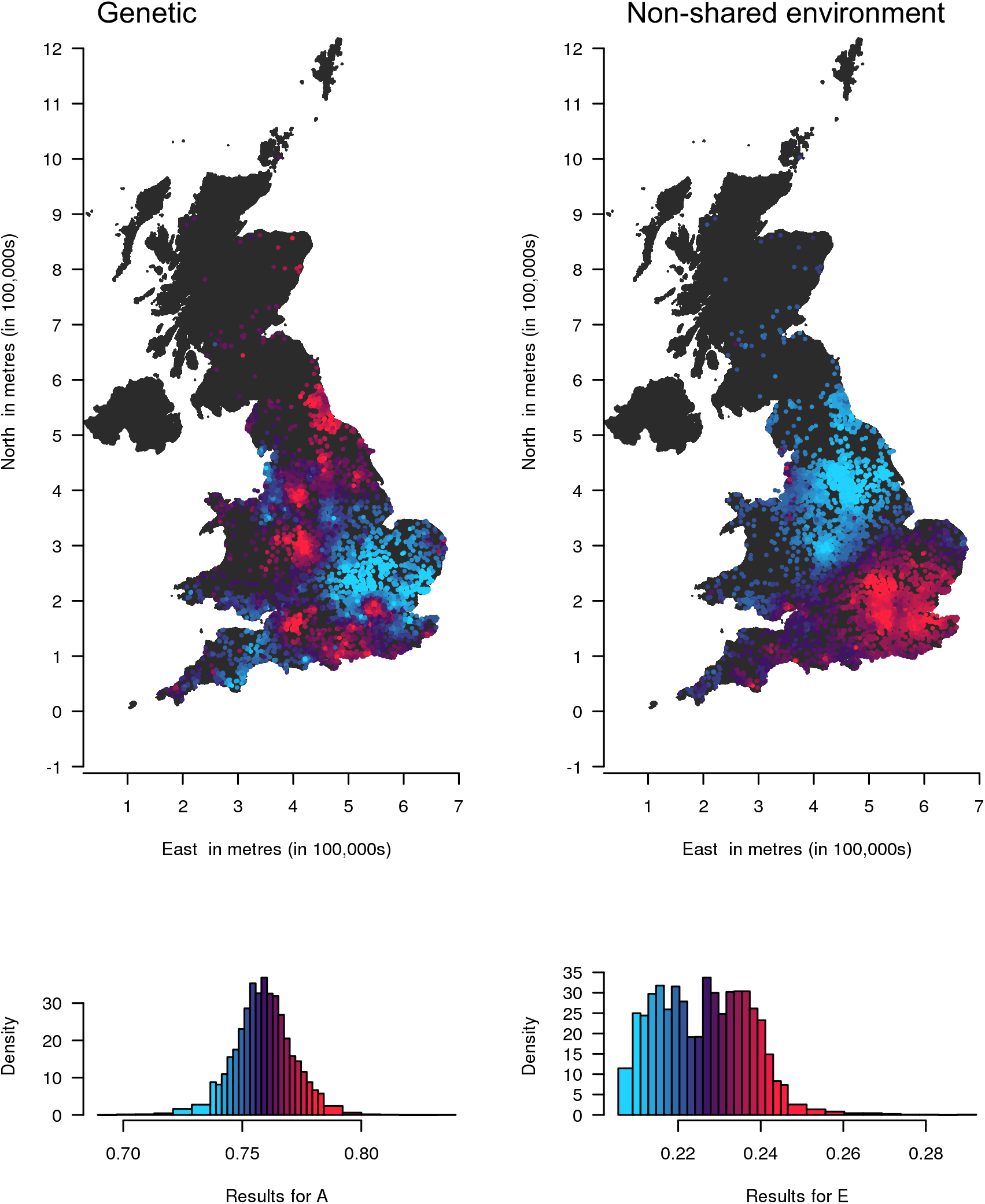
Both genetic and environmental influences on autistic traits in the UK appear to vary geographically across the country. Geographical variation in genetic (A) and non-shared (E) influences on childhood autistic traits in the UK. The contributions of A and E range from low (blue) to high (red). The histograms show the distribution of the estimates, coloured in the same way as the points on the map. The A and E estimates are not standardised and are therefore not constrained to add up to one at each location. A higher-resolution interactive version of this map is available at https://dynamicgenetics.github.io/spACEjs/.

We estimated 95% confidence intervals for A, C and E at each target location and using the CATSS data we performed sensitivity analyses for how A, C and E estimates vary based on the historical residential location used for the twin pairs (**supplementary video 1**). This may allow for identification of critical developmental periods when the geographical environment is particularly influential; for example, if clear patterns are seen when participant locations for the analysis are based on their location at a specific age.

#### Sex limitation models

While some previous studies have identified no aetiological sex differences for ASD (Constantino & Todd, 2003; Ronald et al., 2006), others have. For example, statistically significant but modest quantitative sex differences (i.e. small sex differences in the magnitude of A, C and E influences) were found in previous work with CATSS (Ronald, Larsson, Anckarsäter, & Lichtenstein, 2011), although no sex differences were found in TEDS (Ronald et al., 2006). We examined sex differences in CATSS data and replicated small quantitative sex differences in aetiology (**supplementary Table S2**). To maximise power, we report results that equate the aetiological influences for males and females in the main text, but we include separate analyses for males and females in the supplementary materials (**supplementary Fig S5 and Fig S6**). In practice, these separate analyses produced similar results.

### Data availability

The data used in this study are available to researchers directly from CATSS and TEDS. Procedures for accessing the data are described at https://ki.se/en/meb/the-child-and-adolescent-twin-study-in-sweden-catss and https://www.teds.ac.uk/researchers/teds-data-access-policy.

### Code availability

Code that implements the spACE model and the interactive online maps is available under a GPLv3 open source licence from the scripts directory at https://github.com/DynamicGenetics/spACEjs/.

## Results

### Sample characteristics

Sample characteristics are displayed in Table 1, including the percentage of those scoring above the diagnostic cut-offs used for ASD. Most people in these general population samples score below these cut-offs. The mean scores in our samples compared to those recruited were fairly similar for both the A-TAC in CATSS (0.75 versus 0.82 in the original sample) and the CAST in TEDS (4.95 versus 5.01 in the original sample).

**Table 1.** Sample characteristics for CATSS and TEDS

|  | **CATSS (N=16,677)** | **TEDS (N=11,594)** |
| --- | --- | --- |
| **Age of assessment** | 9 to 12 | Mean=11.30, SD=0.72 |
| **Percentage of males** | 51% | 48% |
| **Median score on autistic traits measurement<sup>1</sup> (IQR)** | 0.00 (0.00 to 1.00) | 4.0 (2.14 to 6.98) |
| **Percentage scoring above cut-offs for autistic traits measures** | Above 4.5 = 3.24%<br>Above 8.5 = 0.94% | Above 15 = 2.22% |

<sup>1</sup>*This was the Autism-Tics, ADHD and other Comorbidities (A-TAC) for the Child and Adolescent Twin Study in Sweden (CATSS) and the Childhood Autism Spectrum Test (CAST) for the Twins Early Development Study (TEDS).*
*SD= standard deviation, IQR=interquartile range*

### Mapping the aetiology of autistic traits in Sweden

We mapped the results from each of the 4,199 locations (**Figure 3** and https://dynamicgenetics.github.io/spACEjs/). Since we modelled raw variance (i.e. we did not standardize the A, C and E estimates at each location to add up to one), genetic and environmental influences are not reciprocal at each location, so it is possible for a location to show both strong genetic and environmental influences. Maps with A and E constrained to add up to one in each location (i.e. proportional) are shown in **supplementary Fig S2** and show similar results to those for the raw variance. For comparison, we have also plotted results of the weighted means of scaled autistic trait scores at the same locations in **supplementary Fig S3** and observe geographical variation for mean autistic trait scores, reflecting the expected variation in prevalence. A map of the unstandardised total variance is shown in **supplementary Fig S4**. While we replicated previous reports of quantitative sex differences in A, C and E components in CATSS at the population level, the differences were small (supplementary Table S2) and separate analyses of males and females produced similar results (supplementary Fig S5 and Fig S6).

The results suggest that the amount of variation in autistic traits explained by genetic infuences (A) appears higher in the far south and in a band running across the centre of the country. Environmental influences appear greatest in the south and north, with reduced environmental influence across the central band.

The histograms for the raw variance indicate that the variance explained by genetic influences ranges from 0.55 to 0.91, with a mean of 0.65 (SD=0.02). The variance explained by E ranges from 0.25 to 0.46, with a mean of 0.34 (SD=0.02). Variation in autistic traits explained by C was approximately zero over the whole of Sweden. Confidence intervals for estimates at each location are provided in **supplementary Table S1. Supplementary video 1** shows that the overall patterns for variation in A and E remained similar irrespective of the historical location used for each twin pair.

### Mapping the aetiology of autistic traits in the UK

**Figure 4** (and https://dynamicgenetics.github.io/spACEjs/) maps genetic and environmental influences on autistic traits in the UK at 6,758 locations. Again, this is a map of the raw variance. Maps with A, C and E constrained to add to one at each location are shown in **supplementary Fig S7**, with very similar results. A map of the unstandardised total variance is also shown in **supplementary Fig S8**.

In the UK genetic influences appear greater in the south, particularly in more central southern areas and the southeast, the Midlands and the north of England. Environmental influences appear greatest in the south and east of the UK, with less influence in the north and the west. The mean of A is slightly higher in the UK (0.76, SD=0.01) than in Sweden (0.65, SD=0.02). Again, C is approximately zero for autistic traits across all regions. As the histograms show, A is fairly normally distributed between 0.69 and 0.84 across regions. E ranges more narrowly from 0.21 to 0.29 in a bimodal distribution with a positive skew. Confidence intervals for estimates at each location are provided in **supplementary Table S3.**

## Discussion

Our results are consistent with previous population-level estimates of genetic and environmental influences, and demonstrate geographical variation in genetic and nonshared environmental influences on autistic traits in Sweden and the UK.

Geographical differences in genetic influences on autistic traits may be indicative of gene-environment interactions where the interacting environmental variable varies by location. For areas of increased genetic influence this means that the environment in these areas draws out genetic influences, in the same way that the presence of airborne pollen would reveal individual differences in genetic risk for hay fever. Alternatively, differences in the A component between areas could reflect active gene-environment correlation, where individuals seek out environments such as cities in part due to their genetic predispositions. It is also plausible that some of the results we observe could be due to geographically patterned genetic differences interacting with genetic and environmental risk factors for autistic traits. We have only included individuals with the same reported ancestry in the analyses to mitigate the effects of broad demographic differences in genetic background between areas linked to patterns of immigration. However, there are still likely to be smaller regional differences between people of the same ancestral background (Leslie et al., 2015). Areas of increased environmental influence imply regions where autistic traits are more affected by environmental variation. By studying these aetiological interactions in a systematic way, rather than constraining ourselves to a specific measured environment, we can develop novel hypotheses about currently unknown environmental influences.

Our findings complement previous research on geographical prevalence differences in ASD (Bakian et al., 2015; Campbell et al., 2011; Chen et al., 2008; Hoffman et al., 2017; Mazumdar et al., 2013). Similarly, alongside aetiological differences, we observe geographical variation in mean autistic trait scores. These mean differences may be linked to aetiological differences. For example, areas of greater prevalence could represent regions where the environment triggers genetic predisposition to autistic traits. This provides a basis for future research into specific geographically distributed environments that draw out or mitigate genetic or environmental risk, which could in turn be useful for population health measures seeking to reduce the impact of ASD.

We hope these results will spark hypotheses about what these factors could be and inspire future studies to identify them. For example, by examining the maps, one might hypothesise that there is generally higher genetic influence in more densely populated areas and lower genetic influence in less populated regions. This may suggest that urban environments draw out genetic differences in predisposition to autistic traits between people of the same ancestral background (rather than implying greater genetic diversity in these regions). These geographically distributed environments might include psychosocial factors such as the stress of urban living or income inequality, or aspects of the physical environment such as air pollution. This explanation fits with neuroscience literature that suggests living in an urban environment is associated with specific neural correlates in response to stress (Lederbogen et al., 2011). However, urban-rural differences may also reflect differences in service availability or special educational needs programs, which could be involved in revealing these geographical genetic differences in autistic traits. For example, in areas where these services are more readily available this may affect the presentation of autistic traits. The literature on prevalence suggests that other potentially important geographical factors may include healthcare accessibility, diagnostic bias, parental awareness, socio-economic status, neighbourhood deprivation, area infrastructure, or green space accessibility. However, factors such as rater effects or access to healthcare are not likely to play a role in this aetiological variation as we used data from structured interviews in population representative samples, and environmental influences on prevalence are not necessarily the same as environmental influences on aetiology. The maps suggest lower non-shared environmental influences in and around Stockholm, the Swedish capital, compared to the rest of the country. Therefore, it may be that there are environments related specifically to living in or around the capital that result in decreased non-shared environmental influences compared to other areas in Sweden. To examine these potential specific risk factors further and explicitly model interactions between specific environments and variance components, one might fit a continuous moderator model, which examines A, C and E influences as a function of a moderator variable.

Whilst we see similarities in patterns of aetiology between Sweden and the UK for autistic traits, there are also substantial differences. There are several possible reasons for this. It could be due to differences in the measurement of autistic traits in the cohorts, or environmental differences between the two countries. For example, differences in the level of awareness of ASD and therefore possible differential reporting in autistic traits, or differences in the physical or social environments, which may vary between countries in the same way as they do within each country. The diagnostic criteria used for ASD have been shown previously to impact prevalence estimates across different countries (J. G. Williams, Higgins, & Brayne, 2005). Prevalence estimates have been shown to vary between the UK and Sweden and also within these countries on a study by study basis (Chiarotti & Venerosi, 2020; Idring et al., 2015). This may reflect differences in case ascertainment or the screening tools used, but may also reflect socio-cultural or other factors as well. It will be important to investigate this in other countries to explore these international similarities and differences further.

When interpreting these results there are a few important points to consider. First, in some areas the effective sample size is lower than others, for example in densely populated areas the proximity of some twin pairs relative to others can weight their influence relatively highly. In over 99% of target locations effective sample sizes are in the thousands for both identical and fraternal twin pairs (i.e., only 0.33% of locations in Sweden and 0.21% in the UK have an effective sample size less than 1000 for identical or fraternal twin pairs), so estimates remain reasonably precise. Second, due to how the weighting in the analyses works, i.e. participants contribute more to analysis the closer they are to the target location, this results in smoothing over the estimates for A, C and E. The amount by which results over the area are smoothed depends on the tuning parameter used in the weighting. There is a trade-off when selecting the tuning parameter between smoothing over noise and detecting real variation, or between accurately estimating variance components and accurately localising them. Here, we have chosen the tuning parameter to result in some smoothing towards the population mean, but this may mean that some larger localised variation remains undetected. In interpreting the maps, it is important to take into account both the pattern of results shown on the map, and the range of estimates shown by the histogram, while bearing in mind that the effect sizes are smoothed towards the population mean. Third, in common with the previous literature, we find that an ADE model is often a slightly better fit to the data, but here we have not modelled the D component because the high correlation between A and D brings noise to spatial analysis due to switching between the two across locations (Rietveld, Posthuma, Dolan, & Boomsma, 2003). Instead, we interpret A here as a broad genetic component, without the usual connotation of additivity. However, we have included ADE maps in supplementary Fig S9 and Fig S10 for Sweden and the UK, respectively, as these maybe useful for reference.

Fourth, as with any statistical analysis, it is important to consider the assumptions of the model. For twin modelling, these include random mating within the population, that MZ and DZ twins share their environments to the same extent (at least where those environments are not genetically influenced), and that twins are representative of the general population for the traits studied (Rijsdijk & Sham, 2002). These assumptions have generally been found to be reasonable (Evans & Martin, 2000), although there is some evidence to suggest that there is assortative mating for ASD (Nordsletten et al., 2016). This would have the effect of inflating the shared environmental influences, which we find to be approximately zero across locations. For our geographical analyses we do not assume that there is no gene-environment interaction or correlation, because we are explicitly modelling them as our main point of interest.

Fifth, the symptom measures used here may index some symptoms that are common to other traits such as ADHD. However, in the article where we first described the spACE approach (Davis et al., 2012), we examined a wide range of different measures and each gave different results, suggesting that these findings are specific to autistic traits. Although we examine autistic traits in middle childhood in this study, it would be interesting to examine how this aetiology might change across age. Future studies may examine this by incorporating a longitudinal model into the spACE framework.

Sixth, it is likely that, alongside real differences, the patterns include some stochastic variation. Simulation analyses in the original spACE article (Davis et al, 2012) demonstrated that the larger regional differences this approach detects are beyond what would be expected by chance. Despite this, each map will include some stochastic variation. As usual when interpreting the results of any statistical analysis, when interpreting the maps it is important to take account of the effect size by cross referencing the colours from the accompanying histogram. We have calculated confidence intervals for each variance component in each location, and these are included in the supplementary data to facilitate further analysis of targeted hypotheses and specific regional differences.

## Conclusion

Our systematic analysis shows geographical variation in genetic and non-shared environmental influences for autistic traits in both Sweden and the UK. The results will inform further studies of a wider range of measured geographically distributed environments, beyond those already identified as influencing prevalence in the literature. Through these, we will gain greater understanding of how these specific environments draw out or mask genetic predisposition and interact with other environmental influences in the development of autistic traits.

## Acknowledgements

We gratefully acknowledge the ongoing contribution of the participants in the TEDS and CATSS and their research teams.

## Key points

- The prevalence of autistic traits varies geographically, but it was previously unknown whether the underlying aetiological influences vary geographically as well.
- Results from our study suggest that there is geographical variation in genetic and non-shared environmental influences on autistic traits in both the United Kingdom and Sweden.
- Our results may aid identification of previously unknown environmental influences on the aetiology of autistic traits.

