## Supplementary materials for "Mapping the genetic and environmental aetiology of autistic traits in Sweden and the United Kingdom"

*Detailed description of Swedish geography*

There are approximately 9,200 SAMS in Sweden, subdivisions of 290 municipalities. The average population within each SAMS is 1,000 people and therefore the area covered by each SAMS varies by population density.

Overall Sweden is split into a more rural north and central area, known as the lowlands and the more populated areas in a belt from Gothenburg in the west to Stockholm in the east and the very south near Malmö. Much of Sweden is covered by forest and lakes. The capital, Stockholm, is in the east and surrounded by water, with 14 islands in the main part of the city and an additional 24,000 islands it’s archipelago. The archipelago islands are much more sparsely populated and generally provide areas for leisure away from the city. Central Stockholm has a number of key areas, with the old town in Gamla Stan in the South, where parliament is found, with the island of Skeppsholmen in the east and both areas are popular with tourists. The compact city centre, Norrmalm is more of a consumerist area, whereas Kungsholmen in the west is more residential on the edge of Lake Mälaren. Östermalm in the east is a relatively affluent area and Södermalm in the south is historically more working class. Lake Mälaren dominates the countryside west of Stockholm and here the 6^th^ largest city of Västerås can be found, a mix of old and new areas and home to a large technology company.

Gothenburg, in the east is the second largest city and is home the largest port in Sweden and has a rich maritime history. Like, Stockholm, Gothenburg has an old town in the centre, with an area known as the Avenyn in the south, dominated by restaurants, bars, museums, galleries and other leisure spaces. In the south west lies the area of Haga, the oldest working-class suburb in Gothenburg and cosmopolitan Linné is to the west. Similar to Stockholm, although not as vast, Gothenburg also has an archipelago, mostly for leisure but also commuting. Further inland in the area of Värmland, is the largest lake in the country, lake Vänern with the town of Karlstad on its shore. The Värmland area has many areas of forest, farmland and rivers and in the north, wild bears, wolves and lynx can be found and it is much more rural.

The south west of Sweden is a coastal region with fairly low-lying land and many towns and villages, making it the third most populated area in Sweden. As a result, transport links in this area are good and there is even a bridge to Copenhagen in Denmark. Historically this area was fought for between Sweden and Denmark and as a result there are many Danish influences still found today. The main city in this area is Malmö, previously a centre for the shipping industry and as a result is quite industrialised in places. Due to its location, a substantial proportion of the population, particularly in the South, are of non-Swedish ancestry and as such the area is considered very multicultural. The area also has a strong history associated with the Social Democratic party.

Lake Vättern separates the south west from the south east. This area is heavily forested, although glass factories are located within these areas, known as the ‘Glass Kingdom’. Örebro is one of the main towns in the north, on the shore of another large lake and developed due to it being on the way from the south west to Stockholm. Norrköpng is a town where many old mills can be found, as well as Sweden’s largest zoo and another nearby archipelago. The two largest islands in Sweden are also found in this area, Öland and Gotland, both are popular Summer destinations, due to their warmer climate. Öland is linked to the mainland via a bridge and has many forests, meadows and villages dotted around. Gotland is considered a holiday destination, particularly Visby in the Summer, where festivals take place. The rest of Gotland is however mostly countryside and smaller towns/villages.

The northern coastline, known as the Bothnian coast is the most populated area in the north, although it is still considered much less inhabited than the southern areas. The coastline goes all the way up to Haparanda, which is on the Finnish border and has joined with the neighbouring Finnish Tornio to create a borderless ‘Eurocity’ with aspects of both cultures. The coastline consists of cliffs, fjords and some islands with a number of towns along the coast. The largest of these is the University town of Umeå, with a tenth of the Norrland (northern) population living here.

Central Sweden is a sparsely populated, rural, lowland area covered in forests, with numerous lakes and mountains along the Norwegian border. The area is split into 3 regions. Dalarna in the south has many ski resorts and the area of Falun, previously an old copper mining area. The biggest town here is Mora where the inland railway starts and proceeds north, cutting through central Sweden up to the Arctic circle. North of Darlarna is Härjedalen, a scenic area, previously belonging to Norway and thus has many Norwegian influences. The biggest town here is Sveg with a population of around 2,600. Finally, in the north is Jämtland which also has Norwegian influences, as well as the large lake of Storsjön and ski resorts.

Further north of central Sweden is Swedish Lapland, where half of the area is within the Arctic circle. It is a very remote area with many hills, mountains, ancient forests and old mining areas, although a number of small towns and villages are dotted around, and the inland railway also continues and ends in this area. There are many national parks and it is a popular destination for hiking.

*Detailed description of UK geography*

There are over 1.5 million postcode units in the UK, covering, on average, 15 properties. As with SAMS, the area covered by each postcode varies depending on population density.

The United Kingdom (UK) is split into the countries of England, Wales, Scotland and Northern Ireland (although we do not describe Northern Ireland here because it was not included in the TEDS recruitment area and few participants have moved there since recruitment). Generally, the south of the UK has a milder climate compared to the north and has more low-lying land. The UK is a mix of some very urban, previously (or still) industrial areas and more rural traditional countryside areas.

London, a diverse, multicultural city in the south-east, is the capital, with its own distinct boroughs. The River Thames bisects the city with most tourist attractions on the northern half. The city centre consists of the political Westminster area, Buckingham Palace (where the Queen resides), the affluent areas of Mayfair and Marylebone, lively Soho and university areas in Bloomsbury. To the east is the financial hub as well as St. Pauls cathedral. Along the river are Southbank and Southwark, home to many tourist attractions. In the northern area of the city are many affluent residential areas as well gardens, museums and the zoo. West London is generally an area of leisure, shopping and more affluent suburbs extending out towards Heathrow Airport, whereas south London is more residential.

The south-east of England includes the counties of Kent and Sussex which contain many commuter areas. This area is surrounded coastline and linked to Europe by sea, road and rail and its close proximity means the area was historically an area of invasion and therefore a number of castles can be found here. North Kent lies on the Thames Estuary and is very accessible to London, although fairly industrialised. The coastal towns of Kent are often characterised by medieval architecture and maritime history. The town of Dover is home to the busiest ferry port in Europe but is also known for its castle and white cliffs. Kent is also home to Canterbury with its well-known cathedral, head of the Church of England. The High Weald includes parts of Kent and Sussex and is characterised by rolling hills and villages, as well as a number of castles and the spa town of Royal Tunbridge Wells. Sussex has many seaside resort towns ranging from quieter Rye and Eastbourne to livelier Hastings and Brighton.

The south of the UK is fairly rural, with many historical sites, containing the counties of Hampshire, Dorset and Wiltshire, historically known as Wessex. On the coast are the maritime areas of Portsmouth and Southampton. Portsmouth is quite industrialised and very urbanised due to rebuilding after World War 2, but it also has ferry links to the small, scenic Isle of Wight, a popular holiday destination. The New Forest, in this area, is a 220 square mile rural area of forest and coastal areas with a number of small towns and villages. It is an area of ancient laws with free roaming ponies, deer and cows. Dorset has a number of popular seaside resorts including Bournemouth, Weymouth with the Isle of Portland and Lyme Regis. This region is part of the Jurassic Coast, a world heritage site with fossil filled cliffs and also part of the Dorset Coastal path. Chesil beach is also found here, an 18-mile bank of pebble beach. Wiltshire is known for its Neolithic sites of Stonehenge, near Salisbury and Avebury.

More inland are the areas of Oxfordshire and the Cotswolds, where the University city of Oxford is found, a mix of old and new university buildings and a popular tourist area. Oxfordshire is also home to Blenheim Palace, picturesque market towns like Henley-on-Thames and Windsor with Windsor Castle. The Vale of White Horse in south-west Oxfordshire is a rural area with lots of villages and walking routes such as the ancient Ridgeway National Trail. The Cotswolds is a rural area consisting of old, traditional English villages with thatched cottages and locally sourced limestone buildings and a popular destination in the Summer. The larger towns of Cheltenham and Gloucester are found on the western edge of the Cotswolds.

In the west are the counties of Bristol, Bath and Somerset. Bristol is a cosmopolitan, cultural and varied University city on the River Avon. Bristol has different urban and rural areas in and around the city. Bath is also a university city, but also a spa city with old Roman baths from hot springs. Somerset is fairly rural and contains the Mendips, an area of ancient woodland, the Mendip hills, Wookey hole caves and Cheddar Gorge and it is a popular walking area. Somerset is home to places such as the small cathedral city of Wells, Glastonbury, known for its annual festival and the county town of Taunton. The Quantock hills area also located here, with small villages, woodland and streams. Exmoor National Park is another rural site, on the Bristol channel. It is a large area of moorland, woodland and home to diverse wildlife and it extends to the Devon border.

South-west England consists of pre-industrial Devon and Cornwall, popular Summer destinations with many seaside and fishing towns and plenty of farmland. Devon’s main town is Exeter which is the commercial and cultural hub and is well connected. Much of the south coast is known as the English Riviera with old seaside towns such as Torquay and other fishing and port towns. Plymouth city is also found here, historically a naval base but more urbanised today. North Devon also has a number of resort and market towns, whilst central Devon has the large 368 square mile Dartmoor National Park, consisting of moorland, bog land, forest and a number of villages and market towns around it. Cornwall has many harbour towns and is known historically for its china clay deposits. The Eden project, biomes of plants and a major tourist attraction, is located on an old clay pit. Truro is Cornwall’s capital and is a mix of old and new and the northern towns such as Newquay and Padstow and known for surfing. In the far west are the Lizard and Penwith Peninsulas.

East Anglia is an area of flatland, wetlands and coastal areas, consisting of Suffolk, Norfolk, Cambridgeshire. Essex is a commuter area to London and is home to England’s oldest town, Colchester. The Stour Valley in Essex is an area of countryside, historically known for the weaving trade, with a number of villages and towns. Suffolk is more separated by marsh and woodland and has a relatively unspoilt, although fairly eroded coastline. Norfolk is home to the old Norman city of Norwich, which still retains many historical buildings and is fairly isolated by the Fens for a city. The Fens cover a large area of eastern England and used to be fairly uninhabitable until draining took place in the 17^th^ century, resulting in fertile agricultural land with some remaining wetland areas. Another main geographical feature of Norfolk is the Norfolk Broads, an area of wetlands and diverse wildlife. There is also Blakeney point in the north which has a resident seal population. Cambridgeshire is home to the medieval University town of Cambridge.

The west Midlands are a mix of lowlands and hilly areas with the industrial city of Birmingham (England’s second largest city) and the Peak district. Warwickshire’s main towns are Warwick and Coventry, which both have medieval features and Stratford-upon-Avon, home to Shakespeare, is also found here and the county borders onto the Cotswolds. Worcestershire has areas of low-lying land, specifically the Severn Valley and the Vale of Evesham, hillier parts of the Malverns in the west, rural parts in the south and more industrial areas outside of Birmingham in the north. Worcester in the centre is a mix of old buildings and new developments. Herefordshire located over the Malvern hills is a very agricultural area with the county town of Hereford. In the south is the Forest of Dean. Shropshire is one of the largest but least populated counties in England and is along the England/Wales border. Within this area, in the Severn Valley is the Ironbridge gorge with several small villages, historically an area for iron smelting with factories, but now a unique industrial heritage site. However, much of the rest of the county is fairly rural, particularly the Long Mynd heathland in the south. Derbyshire is fairly industrial, but borders on the Peak district National Park in the north.

The east midlands has a number of large urbanised cities and is an old coal mining area, but rural areas can still be found. It consists of Nottinghamshire, Lincolnshire, Leicestershire and Northamptonshire. Nottinghamshire contains the large city of Nottingham and is fairly rural in the north with a number of old coal mining areas and also the Sherwood forest. Leicestershire contains the modern and multi-cultural city of Leicester with a slightly more rural area in the west. In the east is England’s smallest county, Rutland, with Rutland water, popular for water sports and with nature reserve areas. Northamptonshire has the fairly modern town of Northampton as well as a number of picturesque villages and canals. In Lincolnshire, the small cathedral city of Lincoln is located, as well as the Lincolnshire woods in the east, a chalky area with hills and valleys and also the Gibraltar Point National nature reserve of marshland and the Fens. Skegness, a busy seaside resort is also found here.

North-west England has the large cities of Manchester and Liverpool and the seaside resort of Blackpool, but also the unspoilt, mountainous Isle of Man. The county of Cheshire in this area is home to Chester with a mix of medieval, Roman, Tudor and Victorian buildings and with a Roman amphitheatre. In Merseyside, to the west, is Liverpool, an urbanised, historically maritime city with a ferry port that links to the Isle of Man. Finally, in this area is Lancashire, home to Blackpool.

The scenic Lake district, in Cumbria, is also found in the north as well as the varied area of Yorkshire, with urbanised, coastal and rural areas. The Lake District has 16 major lakes, mountains and small towns. The area is popular for holidays, walking and boat trips. To the north of the lake district is Carlisle. Yorkshire has a variety of different areas, from urbanised areas such as the previously industrialised Sheffield city, known for its steel industry, Leeds, considered to be the commercial capital of the area and the working town of Bradford to rural areas like the large Yorkshire Dales National Park and the North York Moors National Park. There are also market and spa towns as well as coastal towns. York, the capital of the area is a historical city, previously central to religion and politics. There are two coastlines here, the east coast with dunes and mud flats and the north coast with resorts and a rich smuggling history. In the north east of England are the areas of Northumberland, Durham and Tyne and Wear. Durham city has a university and cathedral in its centre, but the outer areas are more urban. Nearby are the Pennine Valleys of Teesdale and Weardale. Tyne and Wear is a fairly industrial area on the North Sea. Whereas, Northumberland is more varied with a large National Park, coastline up to the Scottish border, sandy beaches for low-key holidays and nature reserves such as the Farne islands for seabirds.

Wales is split into the more populated and coastal south, hilly, rural Mid-Wales and mountainous north Wales. The south is where most people live and includes the Wye valley full of medieval towns, the Valleys near the mountainous terrain in the north and previously an area of coal mining and working towns. Cardiff, the capital of Wales is also found in the south, on the waterfront and with leisure areas, a sports stadium, a castle and Cardiff University. The second largest city, Swansea is also found here. Also facing the Bristol channel is the Gower Peninsula consisting of sandstone and limestone with cliffs, seaside towns, marshland and ruins and is popular for both hiking and surfing. Carmarthenshire is a quieter area, whilst southern Pembrokeshire has many coastal areas, popular for holidays and walking, along the Pembrokeshire Coastal path within the National Park. Mid and northern Pembrokeshire cover the most westerly area of Wales and has a more wild, rugged coastline with bays, islands, caves and rocky headlands. It is a popular area for outdoor pursuits and water sports and is connected to Northern Ireland via ferry. Mid-Wales is a much more mountainous, scenic area with many small towns and villages dotted around and reflects traditional Welsh culture. The Brecon Beacons National Park can be found here with high hills, the remote Black mountains, moorland, caves, waterfalls and small villages. The Elan valley is also here and consists of four lakes/reservoirs that used to supply Birmingham with water. Montgomeryshire, in the north, is a rural sparsely populated area and the Cambrian Coast in the west is a mountain backed area with a number of coastal resorts such as Cardigan Bay. Finally, North wales is most known for Snowdonia National Park, a fairly barren, but scenic area of mountain range, with Snowdon being the main one at 3560 feet high. This used to be an area for slate mining and has a number of small villages and two main waterfall areas. In the west of this area is the Llŷn, which is cliff lined, hilly and very remote with many people speaking Welsh as their first language. The island of Anglesey is here and consists mostly of pastoral land. The north coast is more populated with the University town of Bangor and holiday destinations.

Scotland is split into the Highlands in the north and the Lowlands in the south. Southern Scotland is home to the main cities of cosmopolitan and medieval Edinburgh and urban Glasgow and this area has coastal towns, forests and agricultural land. Edinburgh was built on extinct volcanoes and is therefore a very hilly area, although the surrounding area of the Lothians is relatively flat. Within Edinburgh’s centre there is a medieval old town with many tourist attractions, the University and Holyrood park, an area of nature within the city which also contains Arthur’s seat, an 823-foot peak. The new town is still over 200 years old but is much more commercial, professional and business centred. Then there is the area called Leith, a vibrant, cosmopolitan port area. Outside of Edinburgh is the coastal east Lothian area on the Firth of Forth in the north and the more affluent Lammermuir hills to the South. Mid Lothian is a hilly area and west Lothian is a more industrialised, old mining area and considered to be less affluent. The largest city in Scotland, Glasgow, is also in the south on the River Clyde. The area around the city, known as the Clyde, opens gradually out into countryside.

The rest of southern Scotland is split into 3 regions; the Borders, Dumfries and Galloway and Ayrshire. The Cheviot hills lie on the border between England and Scotland leading to the Borders area on the River Tweed. There are many ruins and historical attractions here. The southern part of Dumfries and Galloway is home to the Solway coast, also known as the Scottish Riviera. Dumfries is the largest town in this region and is an old sea port town. Kirkcudbright along the coast has a working harbour. The Galloway hills are located in the north along with the Galloway Forest Park, Britain’s largest forest park, with a varied landscape of mountains, lochs, coast and moorland. The Rhinns of Galloway is a peninsula on the Solway coast with ferries to Ireland. Ayrshire is a hilly and agricultural area in the north.

Central Scotland is more varied with large lochs and forests in the west, rural and industrialised areas and fishing villages in the east and peaks in the north. The River Forth runs through the south-east area, where the old town of Stirling is located, considered to be a smaller version of Edinburgh. In the west is Loch Lomond and the Trossachs National Park, easily reached from Glasgow via the ‘gateway’ town of Balloch and so it is a popular area with tourists. The Trossachs consists of forests, peaks, lochs and small towns. Loch Lomond is large and has 37 islands which are mostly privately owned. In the east is the small region of Fife, surrounded on three sides by water with the Firth of Tay in the north, the Firth of Forth in the south and the North Sea in the east. The north of Fife is fairly rural, whilst the south is semi-industrial and is an old coal mining area, like central Fife as well. On the coast is St. Andrews, a University town with historical buildings and also a popular golf destination. The East Neuk on the coast, has many fishing villages and somewhat of a Flemish influence due to its trading history. Perthshire in the north east is a very agricultural, mostly rural area, although the ancient town of Perth and Scotland’s historical capital is found here, flanked by the north and south Inch parks. To the north is Strath Tay with Loch Tay, an area of countryside and woodland. Further north are the highlands and Rannoch Moor, an uninhabitable and fairly inaccessible area of bog land, high land and lochs.

Argyll in the west is a remote area, transitioning between lowland and highland and with numerous islands. The mainland area has some seaside resorts, such as Victorian Oban in the north which has the largest port in north west Scotland. In the middle of this mainland area is Kilmartin valley and glen with many Bronze age and Neolithic remains and in the south is the Kintyre peninsula, a fairly isolated and rural area with mostly farmland. The islands are part of the inner Hebrides and are also quite varied, with the popular holiday destination of the Isle of Bute, the Christian pilgrimage Isle of Iona, the rocky Isle of Coll with only one village and vast sand dunes, the Isle of Islay, famous for its single malt whiskey production and birdlife, the mountainous Isle of Jura with its uninhabited and fairly inaccessible west coast and large deer population. The most accessible island and possibly the wettest is the Isle of Mull with peaks, pastoral land and fishing port. There is also the small uninhabited Isle of Staffa off its west coast and many other islands in this area.

North east Scotland has a number of industrial cities and port towns, although further north becomes mountainous. The post-industrial city of Dundee is located here, along with Scotland’s third largest city, Aberdeen, lying on the rivers Dee and Don with a working harbour. The area is also considerably wet and windy. Between these two cities lies the Angus coast with a number of seaport and fishing towns. North of Dundee is the agricultural Strathmore region, followed by the Grampian Mountains and the Angus Glens. This area of valleys and peaks is sparsely populated and once snow falls becomes impassable, although ski resorts are found here. In the large region of Aberdeenshire and Moray there are many historical sites and it is split into two main areas, the Hinterlands, an area of farmland and valleys and the rocky coast with fishing villages and beaches. The area of Deeside, also known as Royal Deeside due to its connections with the Royal family has a number of villages as well as Balmoral castle. Speyside, based around the River Spey, is known for its numerous whiskey distilleries. Finally, the Moray coast from Aberdeen to Inverness is wilder with farmland inland and fishing villages by the sea.

The Northern two thirds of Scotland are known as the Highlands, a very remote but unspoilt area, with forests, lochs, mountains and rugged coastline. Inverness, on the River Ness, is the main city here and transport hub. The River Spey extends from the Moray Firth inland to the Cairngorms and Strathspey region. The Strathspey is a forested area known for its salmon fishing and the Cairngorms is Britain’s most extensive mountain range with a variety of wildlife and is popular for hiking, water sports and skiing. The Great Glen is a major Faultline across the Highlands with the UK’s highest peak in the South of Ben Nevis. This area also has four large lochs; Ness, Oich, Lochy and Linnhe, all linked by the Caledonian canal. The west coast of the Highlands has fjord-like sea lochs, beaches, cliffs and mountains, moorland and bog land and is the least populated part of Britain. The north west is particularly remote and receives the fill force of the North Atlantic weather. The north coast, also on the Atlantic has a rugged shoreline with mountains in the west and Lochs and grasslands in the east. The east coast is quite different to the west, with moorland and grassland areas and a strong historical and fishing heritage.

Scotland has a number of island clusters, the Inner and Outer Hebrides in the west and the Orkney and Shetland islands in the north. Skye is the main Inner Hebridean island with two main settlements, the Cullin ridge peaks, the scenic Trotternish peninsula in the north, the Sleat peninsula in the south and the hilly, sparsely populated Isle of Raasay offshore. The other Inner Hebrides are quite small islands. Muck is the smallest and privately owned, Eiss, a 1000-foot plateau of an island is owned by the islanders and Rùm with its Cullin peaks and Canna are owned by national agencies and act as nature reserves. The Outer Hebrides consists of approximately 200 islands, with only some of these inhabited. Lewis is the largest and most populated and is mostly flatland. However, the islands are all quite varied between them in the landscape, some hilly whilst others are flatland. Orkney has around 70 islands and 20,000 people. These islands are areas of farmland with many historical sites, but also wet and windy weather is common. In contrast, the Shetland islands are much less fertile, historically relying on fishing, whaling, naval and merchant services. There are around 300 islands, although only 16 are inhabited. In the south the area is mostly farmland whilst the north is wilder.

*Sex limitation models for autistic traits*

Population-level sex limitation model results for autistic traits are shown in **supplementary TableS2**. We used nested sub-models to test for sex differences in aetiology. The common effects model is the most parsimonious model that adequately fits the data, indicating quantitative, but not qualitative, sex differences. However, because this is a large and well-powered sample, the parameter estimates for males and females that are “significantly” different at an alpha of 0.05 are actually within 1% of each other. Maps for males and females separately are shown in **supplementary FigS4 and FigS5**.

**Supplementary TableS2.** Results for the sex limitation model for autistic traits.

| **Model** | **df** | **AIC** | **BIC** | **-2LL** | **Δdf** | ***P-*value** | **Male** | | | **Female** | | | **r_G_** | **r_C_** |
| --- | --- | --- | --- | --- | --- | --- | --- | --- | --- | --- | --- | --- | --- | --- |
|  |  |  |  |  |  |  | **a^2^m** | **c^2^m** | **e^2^m** | **a^2^f** | **c^2^f** | **e^2^f** |  |  |
| Saturated | 16781 | 25705.62 | -92429.08 | 59267.62 | 8 | 3.52x10^-122^ | NA | NA | NA | NA | NA | NA | NA | NA |
| Full (r_G_ free) | 16789 | 26279.50 | -91911.52 | 59857.50 |  |  | 0.66 (0.63, 0.69) | 4.56x10^-13^ (4.56x10^-13^, 0.01) | 0.34 (0.31, 0.37) | 0.65 (0.57, 0.68) | 2.51x10^-11^ (0.00, 2.51x10^-11^) | 0.35 (0.32, 0.38) | 0.55 | 1.00 |
| Full (r_c_ free) | 16789 | 26280.26 | -91910.76 | 59858.26 |  |  | 0.66 (0.63, 0.69) | 2.36x10^-15^ (1.29x10^-27^, 4.48x10^-14^) | 0.34 (0.31, 0.37) | 0.65 (0.57, 0.68) | 7.26x10^-17^ (1.85x10^-26^, 2.85x10^-13^) | 0.35 (0.32, 0.38) | 0.50 | 0.79 |
| Quantitative differences | 16790 | 26278.26 | -91919.80 | 59858.26 | 1 | 0.38 | 0.66 (0.63, 0.69) | 1.64x10^-13^ (2.70x10^-20^, 1.64x10^-13^) | 0.34 (0.31, 0.37) | 0.65 (0.58, 0.68) | 4.69x10^-12^ (8.09x10^-16^, 6.30x10^-02^) | 0.35 (0.32, 0.38) | 0.50 | 1.00 |
|  | | | | | | | **a^2^** | | **c^2^** | | **e^2^** | |  | |
| Homogeneity | 16793 | 26591.64 | -91627.55 | 60177.64 | 4 | 4.90x10^-68^ | 0.66 (0.64, 0.68) | | 1.53x10^-15^ (6.20x10^-23,^ 3.83x10^-14^) | | 0.34 (0.32, 0.36) | | 0.50 | 1.00 |

The results for the full and nested sex limitation model for A-TAC questionnaire assessed autistic traits are shown. Nested models are compared to the full model based on the -2LL of each model and the significance level is indicated by the *p*-value.

df= Degrees of freedom, AIC= Akaike information criterion, BIC= Bayesian information criterion, -2LL= -2 log likelihood, a^2^= additive genetic estimate (m= for males, f= for females), c^2^= shared environmental estimate, e^2^= non-shared environmental estimate, r_g_= genetic correlation for opposite-sex dizygotic twins, r_c_= shared environmental correlation for opposite-sex dizygotic twins.

**Supplementary FigS1.** Histogram of A-TAC assessed autistic traits in CATSS.

Autistic traits

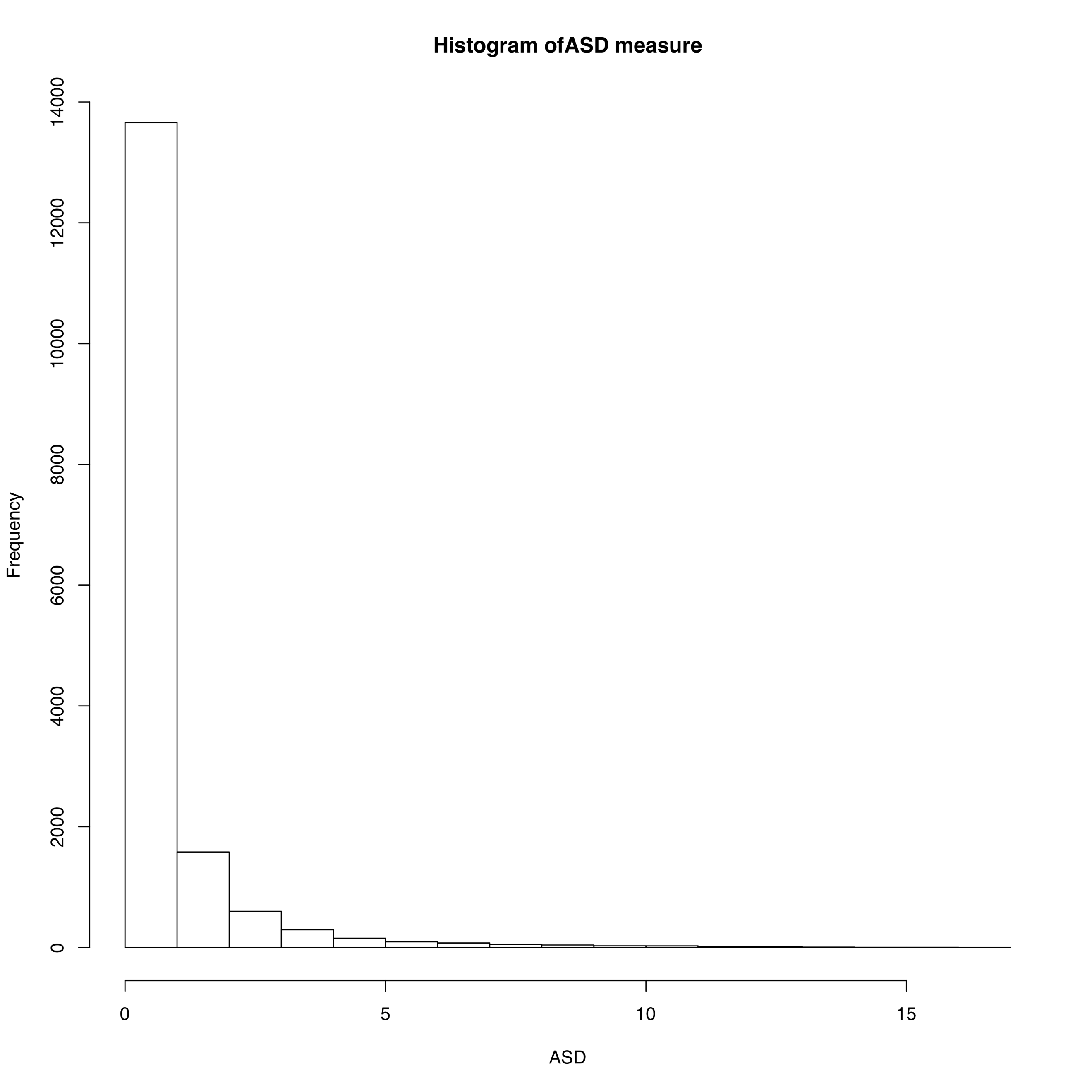

CATSS= Child and Adolescent Twin Study in Sweden

**Supplementary FigS2.** A and E parameter maps and histograms as a proportion of the total variance for autistic traits in Sweden.

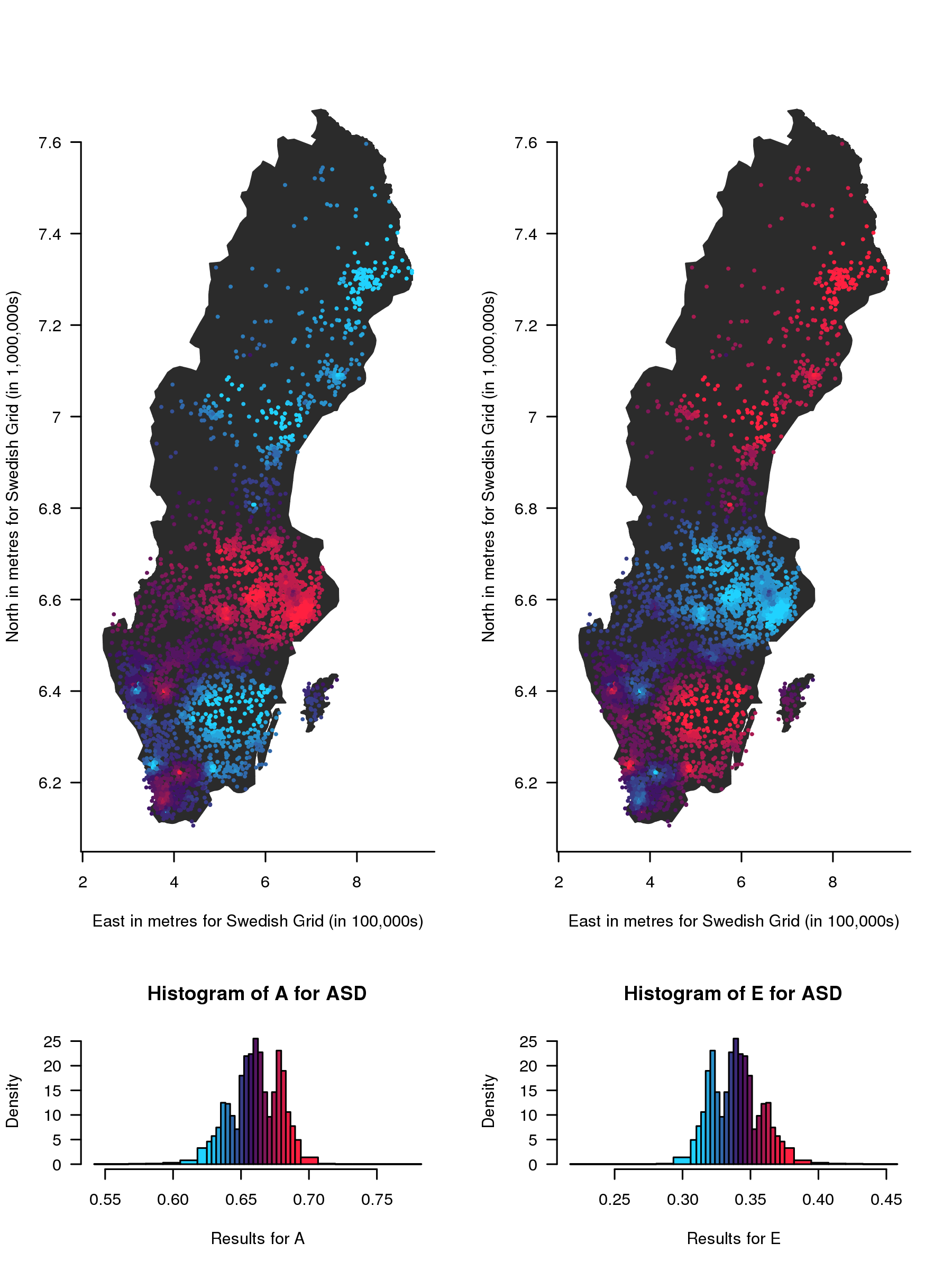

Genetic

Non-shared environment

*Geographical variation in genetic (A) and non-shared environmental (E) influences as a proportion of the total variance on childhood autistic traits in Sweden (results are overlaid on an outline of the SAMS areas). The contributions of A and E range from low (blue) to high (red). The histograms below show the distribution of the estimates, coloured in the same way as the points on the map.*

**Supplementary FigS3.** Weighted mean of scaled autistic traits

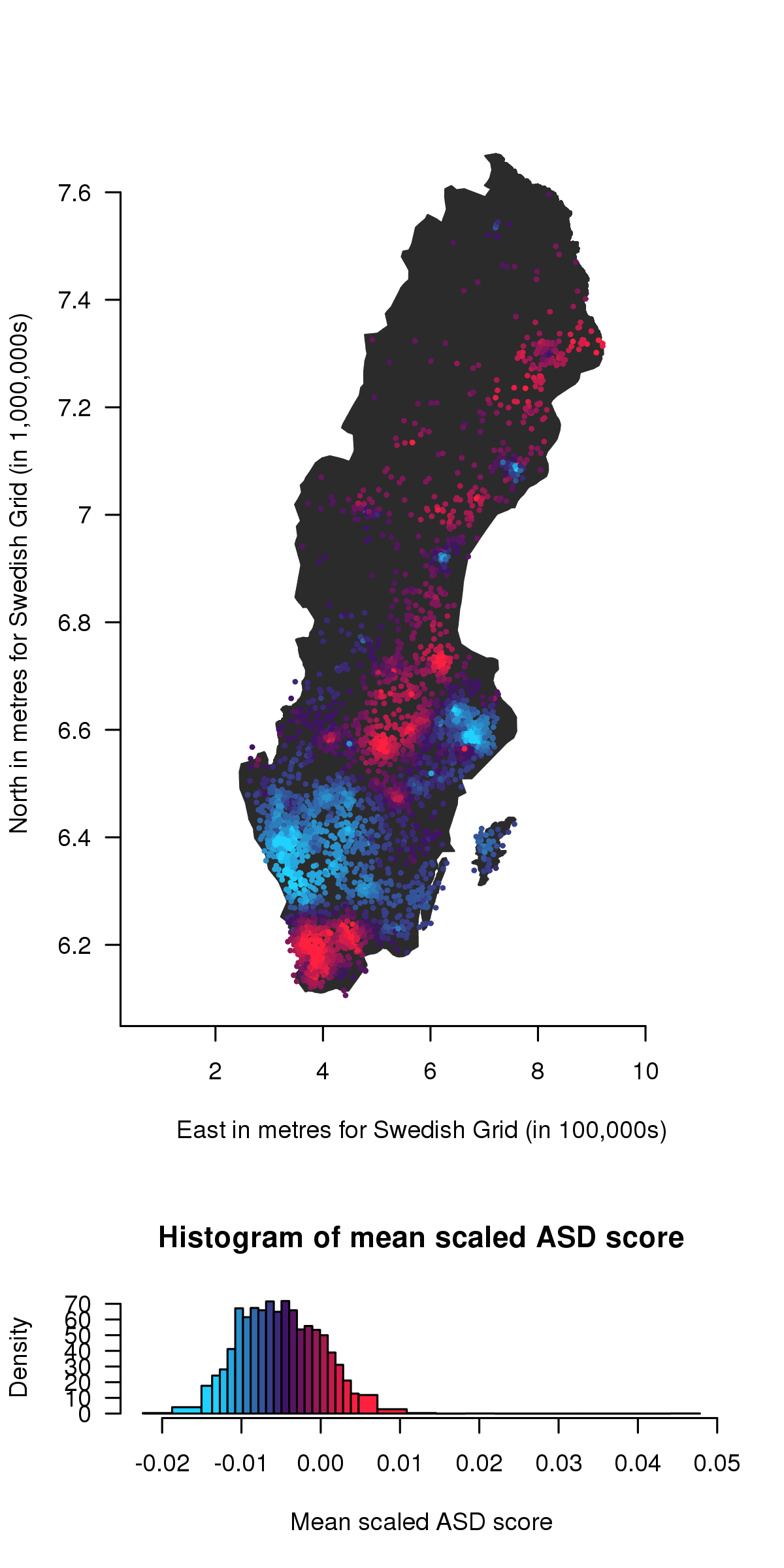

Mean scaled autistic traits

*Weighted mean of scaled autistic traits, from low (blue) to high (red) (these are overlaid on an outline of the SAMS areas). The histograms below show the distribution of the mean scores coloured in the same way as the points on the map.*

**Supplementary FigS4.** Total variance (V) for autistic traits in Sweden

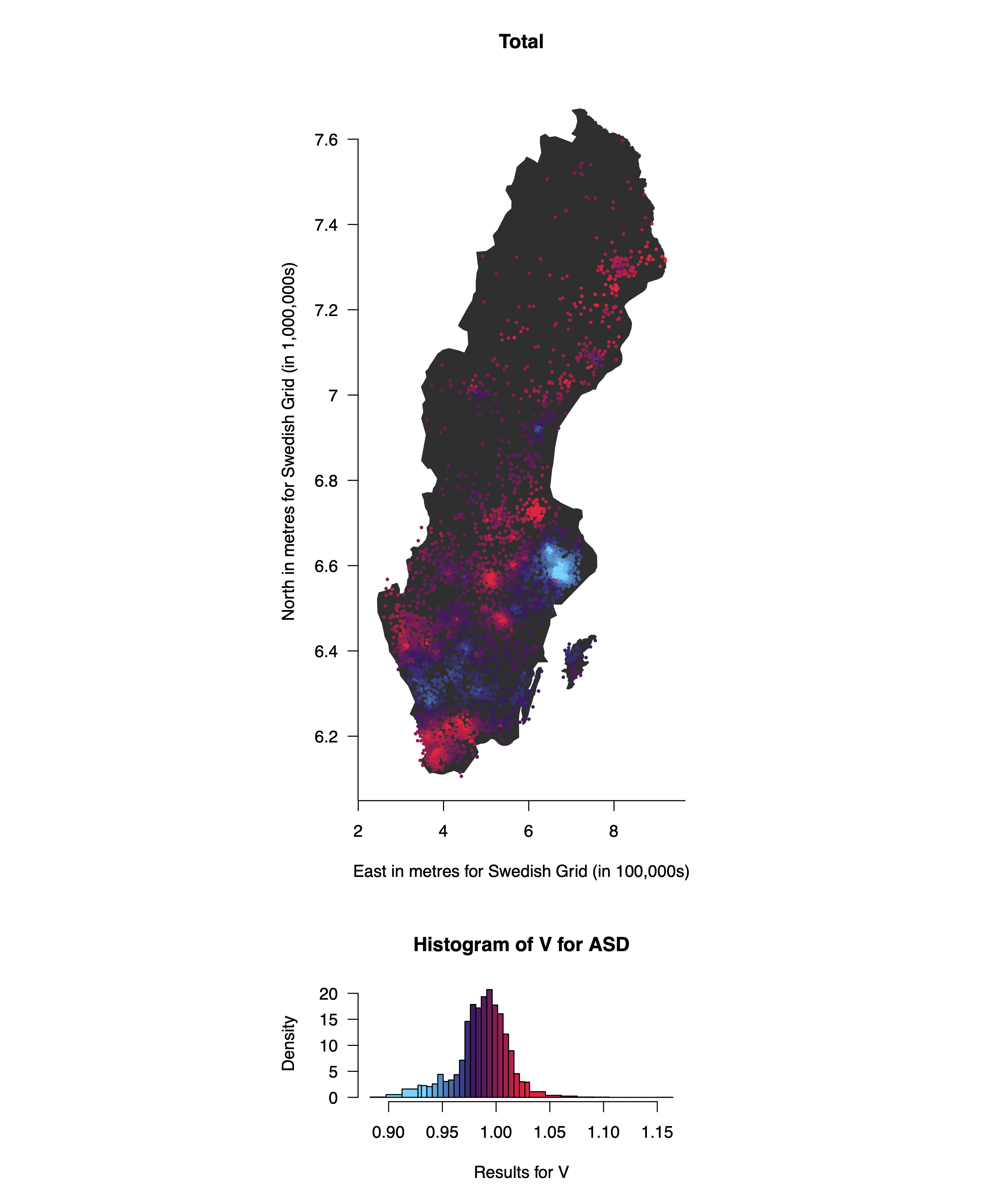

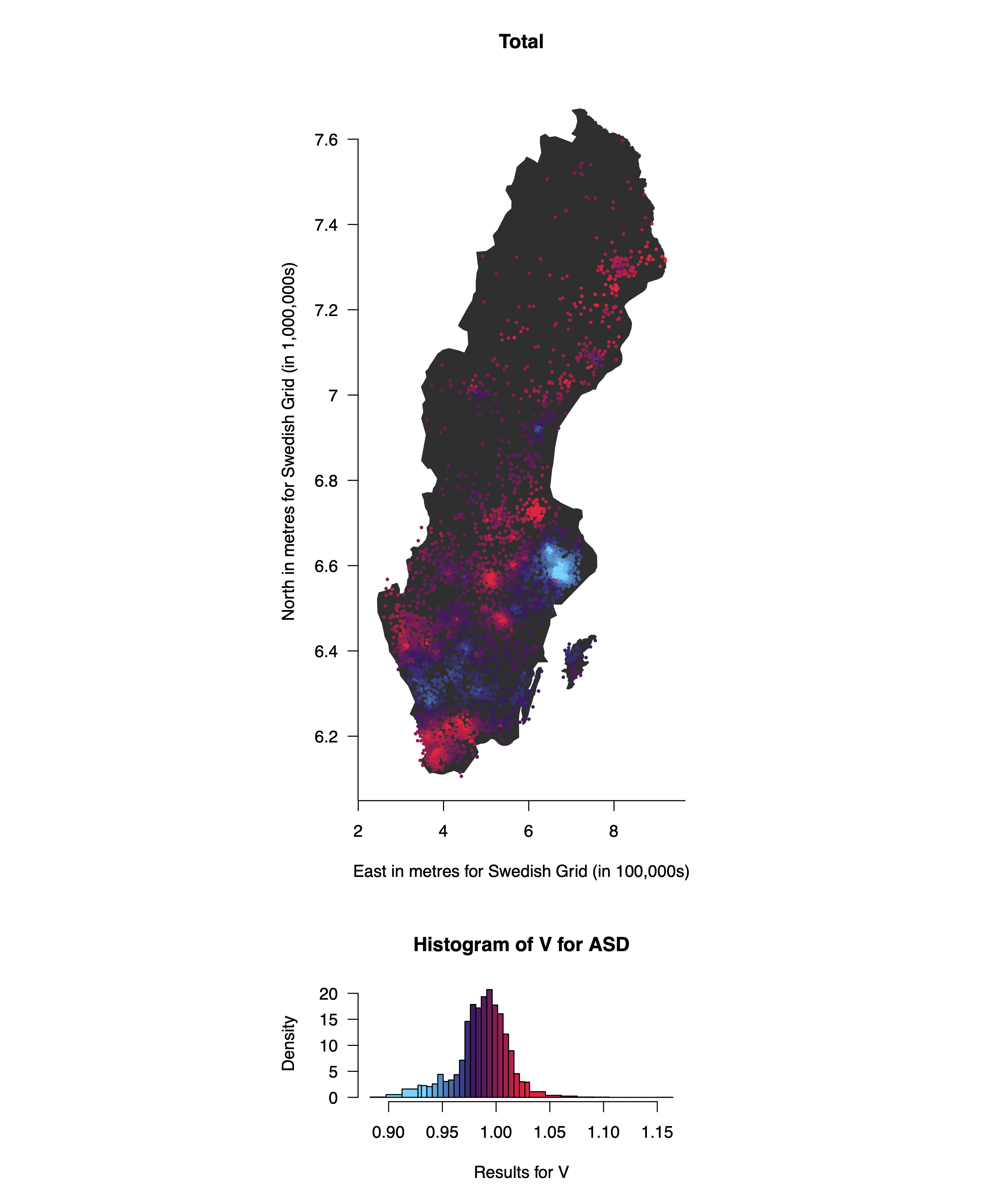

*Total variance for autistic traits, from low (blue) to high (red) (these are overlaid on an outline of the SAMS areas). The histograms below show the distribution of the total variance coloured in the same way as the points on the map.*

**Supplementary FigS5.** Genetic and non-shared environmental influences on autistic traits in males are similar to those overall, although both are lower in the south

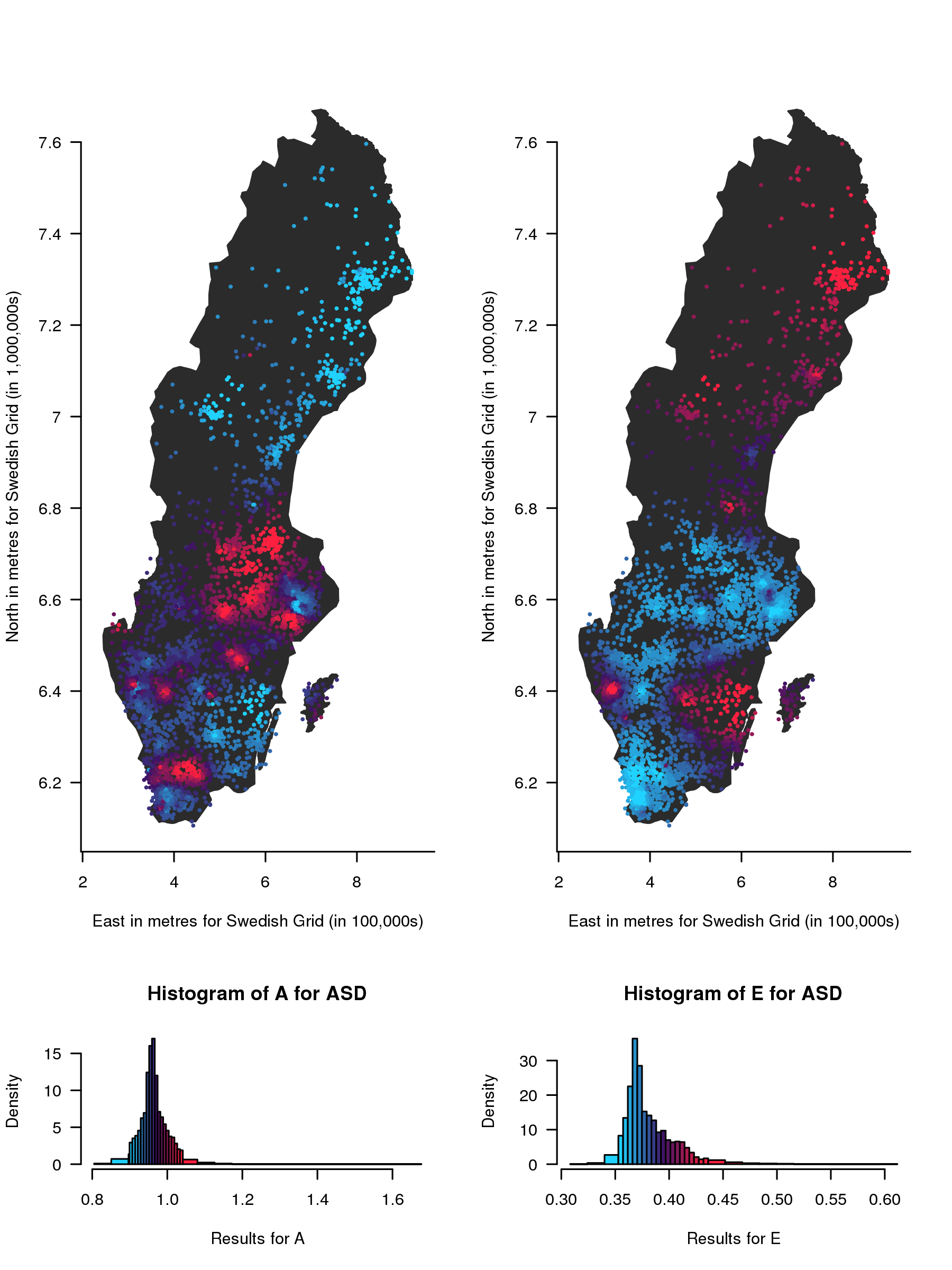

Genetic

Non-shared environment

*Geographical variation in genetic (A) and non-shared environmental (E) influences on childhood autistic traits in Sweden for males only (results are overlaid on an outline of the SAMS areas). The contributions of A and E range from low (blue) to high (red). The histograms below show the distribution of the estimates, coloured in the same way as the points on the map. The estimates are not standardised and are therefore not constrained to add up to one.*

**Supplementary FigS6.** Genetic and non-shared environmental influences on autistic traits in females are similar to those overall, although genetic influences are lower in the south and higher in the north and we also see some evidence for variation in shared environmental influences

*Geographical variation in genetic (A), shared (C) and non-shared environmental (E) influences on childhood autistic traits in Sweden for females only (results are overlaid on an outline of the SAMS areas). The contributions of A and E range from low (blue) to high (red). The histograms below show the distribution of the estimates, coloured in the same way as the points on the map. The estimates are not standardised and are therefore not constrained to add up to one.*

Genetic

Non-shared environment

Shared environment

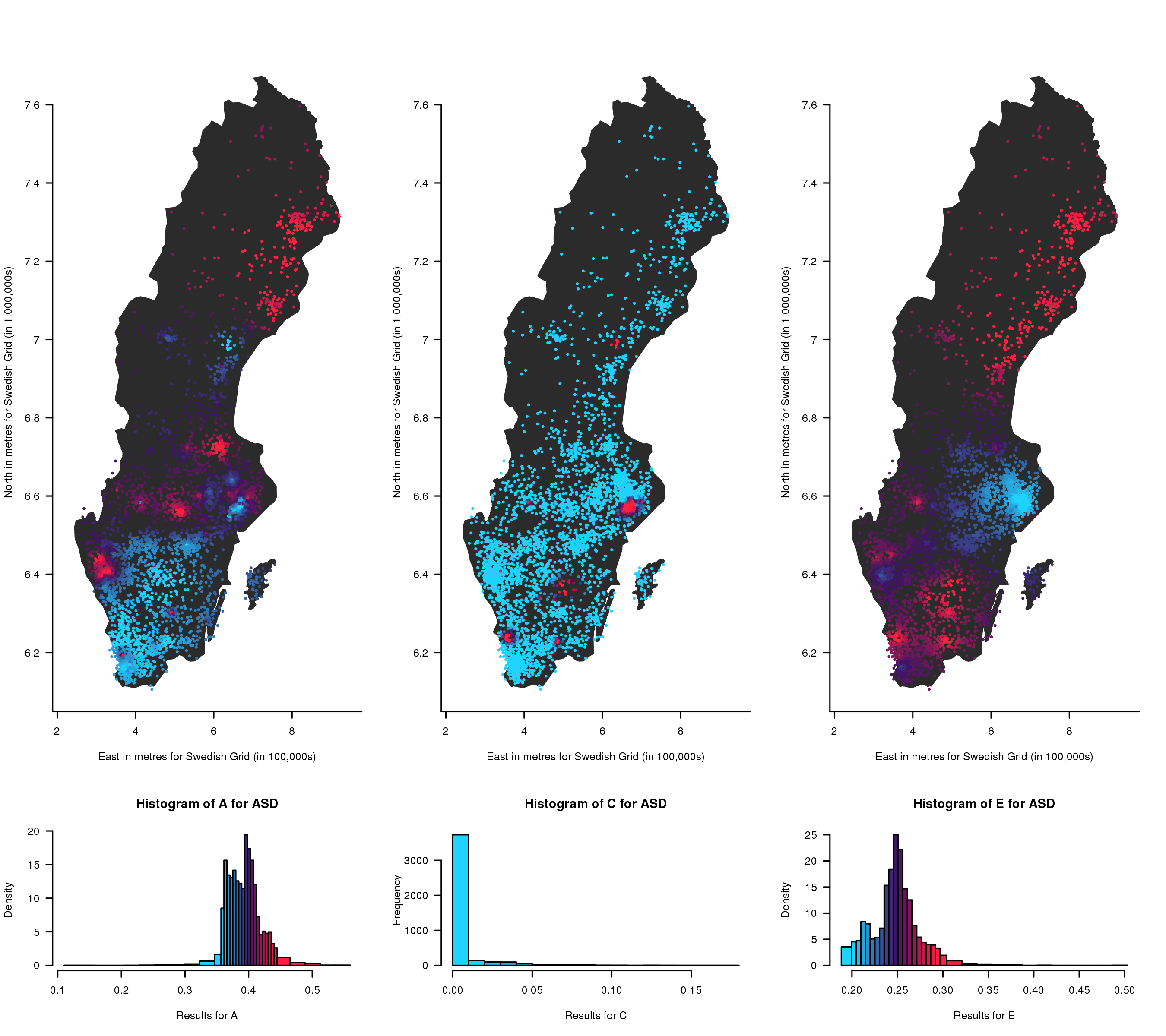

**Supplementary FigS7.** A and E parameter maps and histograms as a proportion of the total variance for autistic traits in the UK

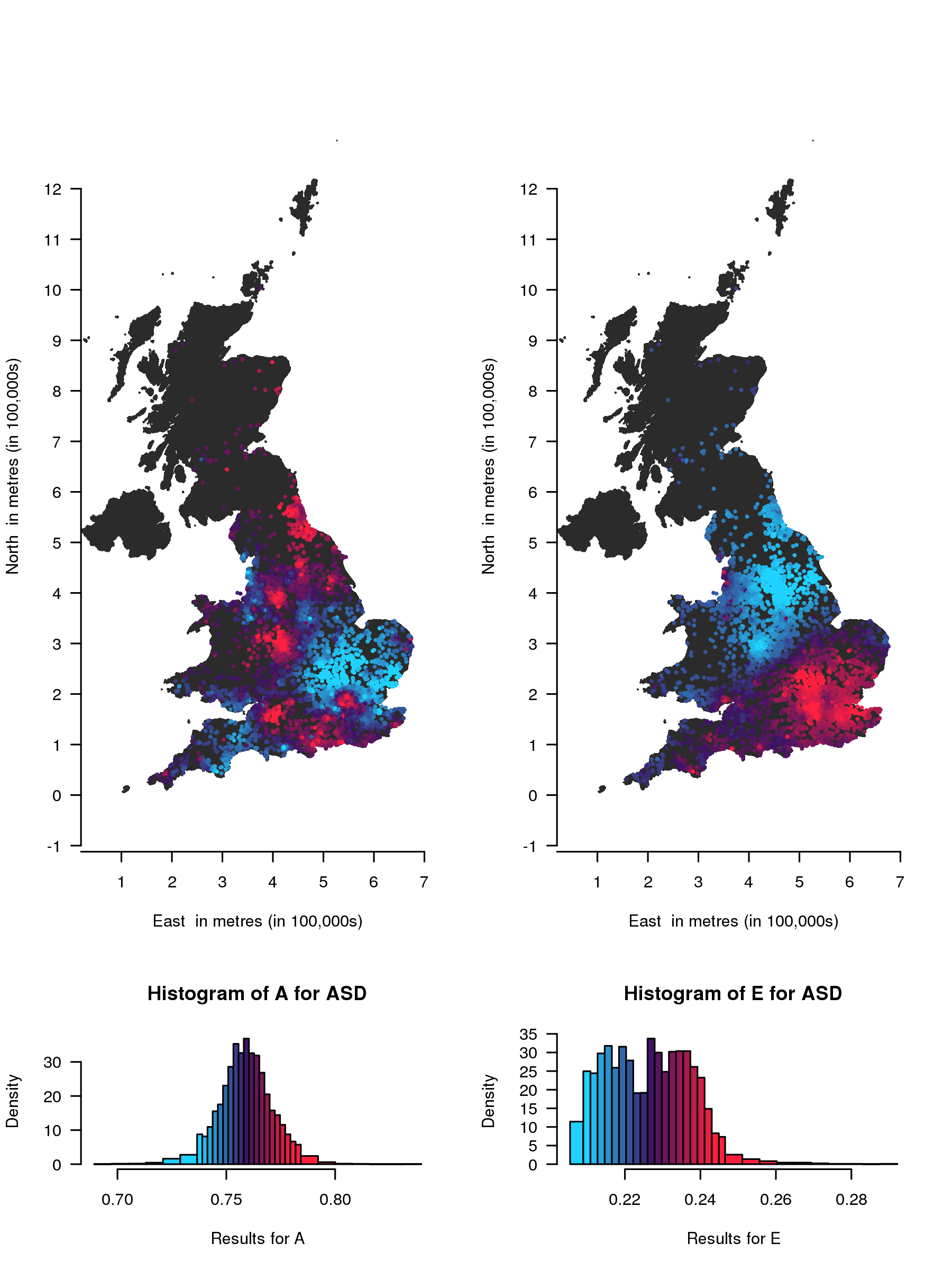

Genetic

Non-shared environment

*Geographical variation in genetic (A) and non-shared environmental (E) influences as a proportion of the total variance on childhood autistic traits in the UK. The contributions of A and E range from low (blue) to high (red). The histograms below show the distribution of the estimates, coloured in the same way as the points on the map.*

**Supplementary FigS8.** Total variance (V) for autistic traits in the UK

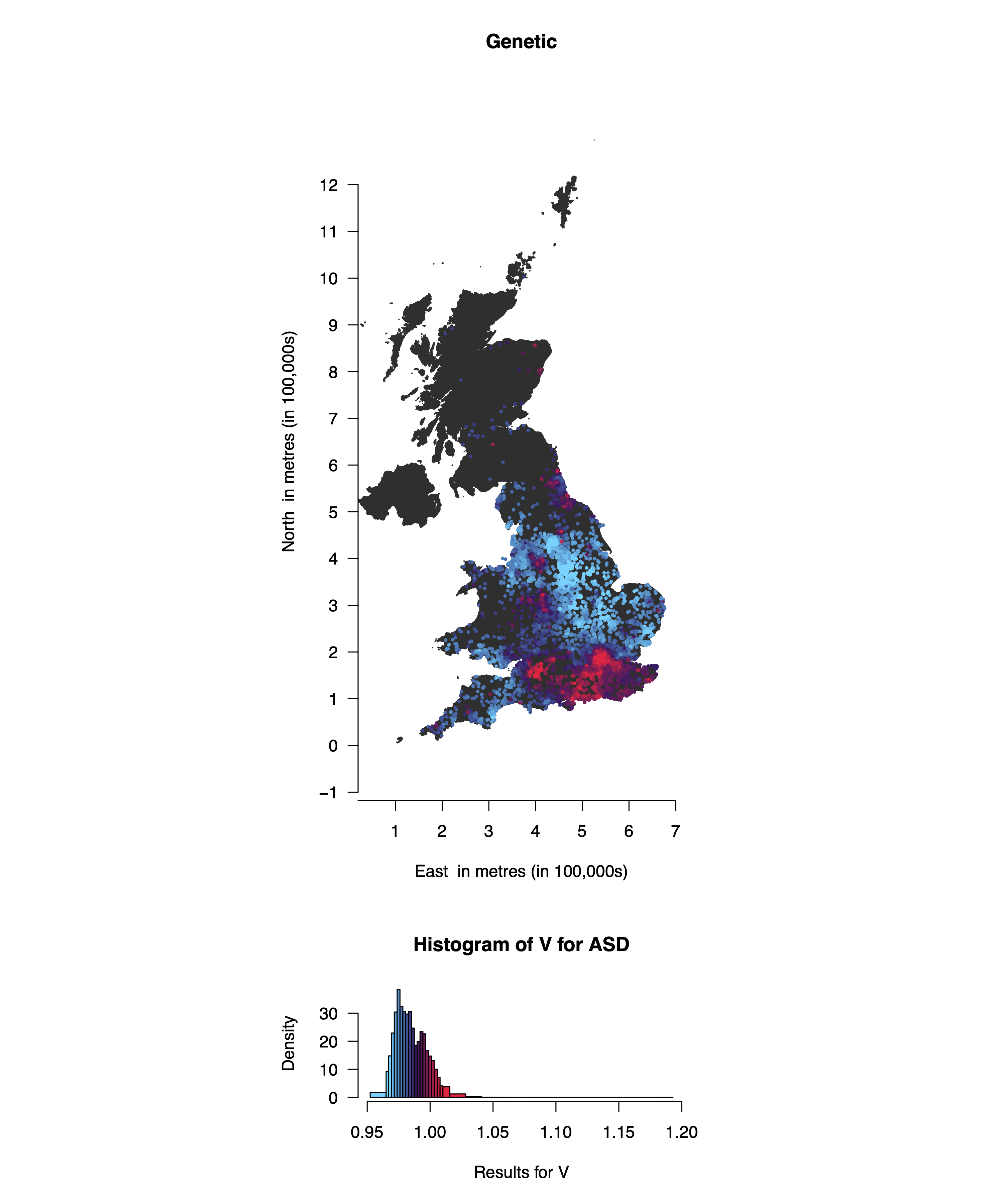

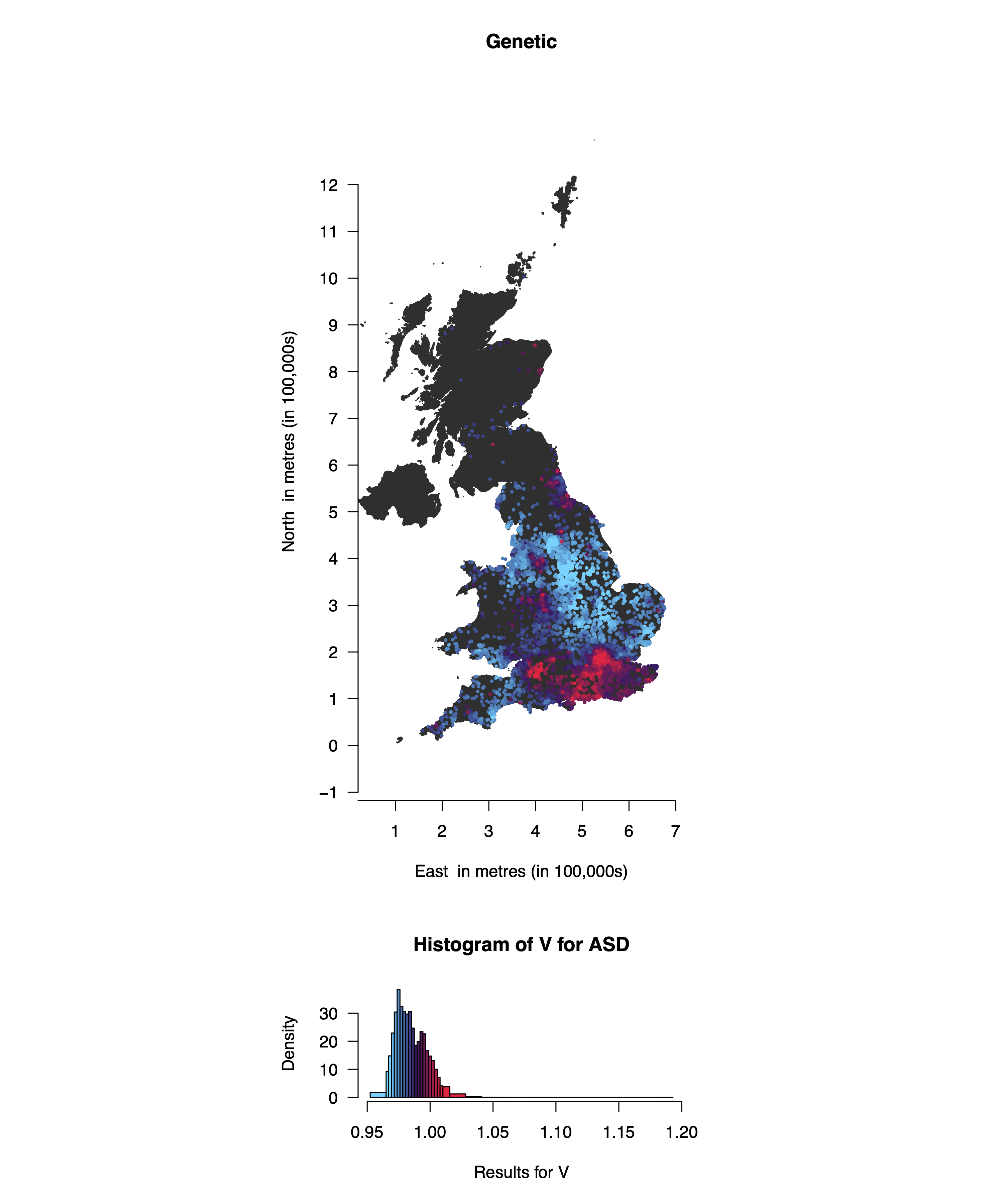

*Total variance for autistic traits, from low (blue) to high (red). The histograms below show the distribution of the total variance coloured in the same way as the points on the map.*

**Supplementary FigS9.** ADE parameter maps and histograms as a proportion of the total variance for autistic traits in Sweden.

*Geographical variation in additive genetic (A), non-additive genetic (D) and non-shared environmental (E) influences on childhood autistic traits in Sweden (results are overlaid on an outline of the SAMS areas). The contributions of A, D and E range from low (blue) to high (red). The histograms below show the distribution of the estimates, coloured in the same way as the points on the map. The estimates are not standardised and are therefore not constrained to add up to one. Higher additive genetic influences are observed in the north and in two central bands from west to east, whereas non-additive genetic influences are higher in the very south and in one central area and north of Stockholm. Non-shared environmental influences are similar to those in the ACE maps other than lower influences in the south.*

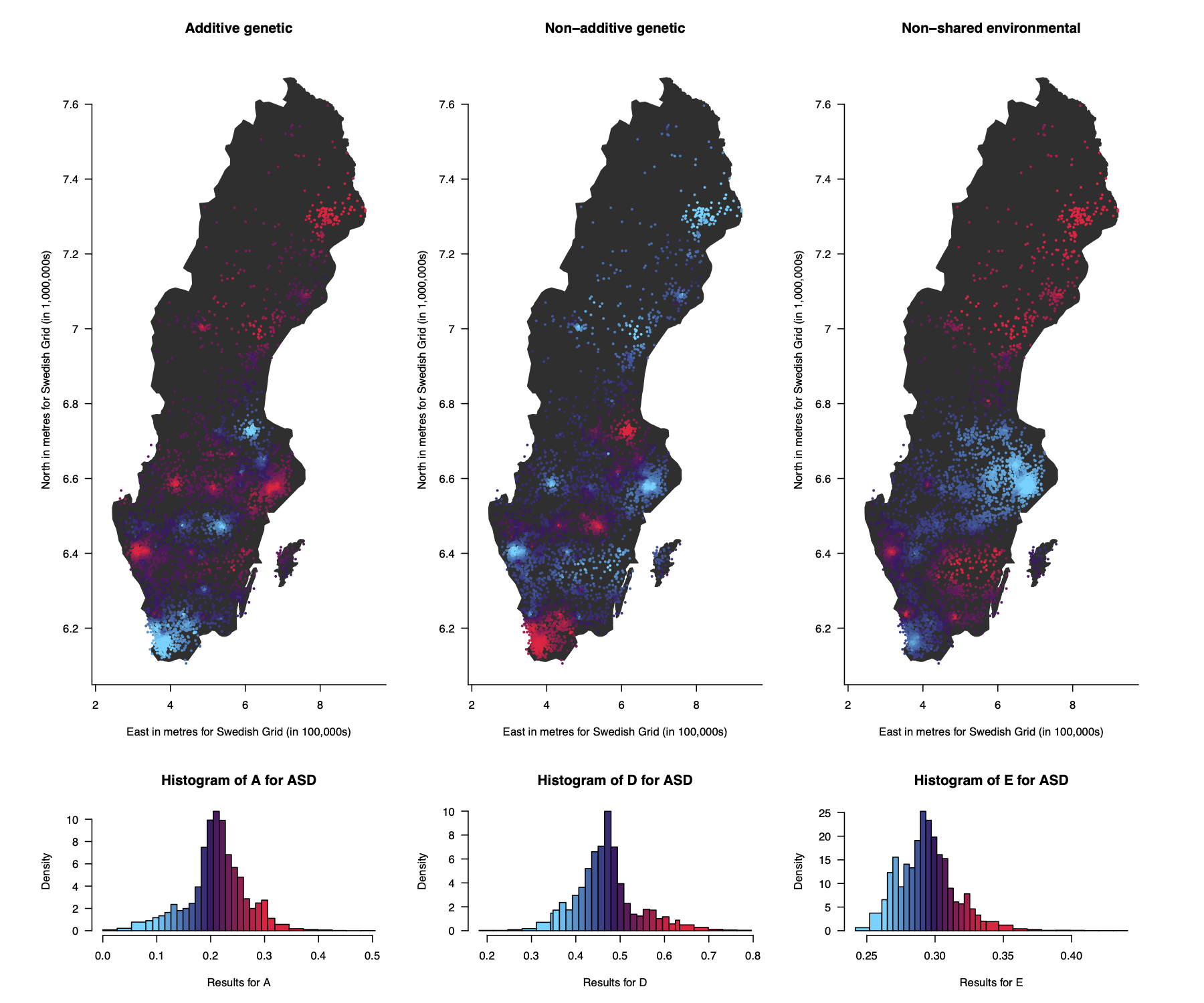

Additive genetic

Non-additive genetic

Non-shared environment

**Supplementary FigS10.** ADE parameter maps and histograms as a proportion of the total variance for autistic traits in the UK.

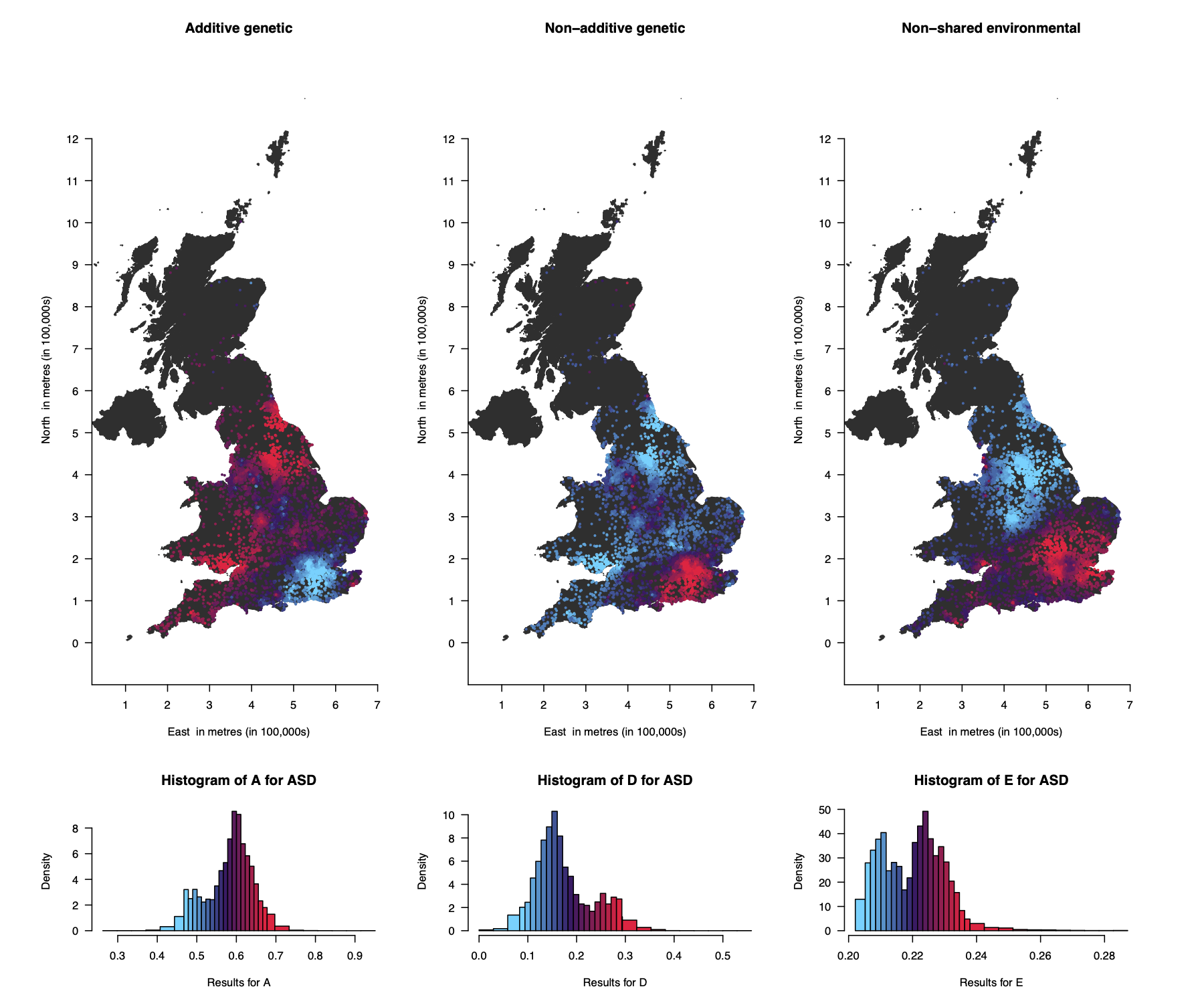

Additive genetic

Non-additive genetic

Non-shared environment

*Geographical variation in additive genetic (A), non-additive genetic (D) and non-shared environmental (E) influences on childhood autistic traits in the UK. The contributions of A, D and E range from low (blue) to high (red). The histograms below show the distribution of the estimates, coloured in the same way as the points on the map. The estimates are not standardised and are therefore not constrained to add up to one. Higher additive genetic influences are observed in most areas other than the majority of the southeast and some central areas. Non-additive genetic influences are higher in the southeast and some central areas. Non-shared environmental influences are similar to those in the ACE maps.*

**Supplementary video 1.** This video shows results from the ACE twin modelling for autistic traits using the twin pairs locations at each year from birth to their 9^th^ year (results are overlaid on an outline of the SAMS areas). To do this we repeated analyses based on participants’ locations at different ages and we combined the resulting maps into a video. The maps are fairly consistent for each year, with slight changes in variation for A and E.
